# Subclonal somatic copy number alterations emerge and dominate in recurrent osteosarcoma

**DOI:** 10.1101/2023.01.05.522765

**Authors:** Michael D. Kinnaman, Simone Zaccaria, Alvin Makohon-Moore, Brian Arnold, Max Levine, Gunes Gundem, Juan E. Arango Ossa, Dominik Glodzik, M. Irene Rodríguez-Sánchez, Nancy Bouvier, Shanita Li, Emily Stockfisch, Marisa Dunigan, Cassidy Cobbs, Umesh Bhanot, Daoqi You, Katelyn Mullen, Jerry Melchor, Michael V. Ortiz, Tara O’Donohue, Emily Slotkin, Leonard H. Wexler, Filemon S. Dela Cruz, Meera Hameed, Julia L. Glade Bender, William D. Tap, Paul A. Meyers, Elli Papaemmanuil, Andrew L. Kung, Christine A Iacobuzio-Donahue

## Abstract

Multiple large-scale tumor genomic profiling efforts have been undertaken in osteosarcoma, however, little is known about the spatial and temporal intratumor heterogeneity and how it may drive treatment resistance. We performed whole-genome sequencing of 37 tumor samples from eight patients with relapsed or refractory osteosarcoma. Each patient had at least one sample from a primary site and a metastatic or relapse site. We identified subclonal copy number alterations in all but one patient. We observed that in five patients, a subclonal copy number clone from the primary tumor emerged and dominated at subsequent relapses. *MYC* gain/amplification was enriched in the treatment-resistant clone in 6 out of 7 patients with more than one clone. Amplifications in other potential driver genes, such as *CCNE1*, *RAD21*, *VEGFA*, and *IGF1R*, were also observed in the resistant copy number clones. Our study sheds light on intratumor heterogeneity and the potential drivers of treatment resistance in osteosarcoma.

**Significance:** Subclonal copy number clones emerged and dominated in relapsed osteosarcoma, with *MYC* gain/amplification being the defining characteristic in our cohort. Selective pressure from neoadjuvant chemotherapy revealed this clone at the time of primary resection, highlighting that genomic profiling at this time may identify clones that are selected for, or determine innate resistance to primary chemotherapy.

## Introduction

Osteosarcoma is an aggressive bone tumor which primarily affects children and young adults. Patients who present with metastatic disease at diagnosis have a poor overall prognosis and those with an inferior response to neoadjuvant chemotherapy have a high risk for recurrence^1–3^. Multiple large-scale tumor genomic profiling efforts have been undertaken to describe the genomic underpinnings and identify new potential therapeutic targets for osteosarcoma^4–9^. These studies revealed that osteosarcoma, typically characterized by a high degree of chromosomal instability, has a large number of chromosomal deletions, translocations, and amplifications. The common alterations present in osteosarcoma primarily involve tumor suppressor genes (e.g., *TP53*, *RB1*, *PTEN*), whereas targetable activating mutations are rare, making it challenging to link the mutational genotype to a broadly applicable treatment strategy^10, 11^. However, recent studies have suggested that targeting focal gene amplifications in consensus driver genes may be an effective strategy for identifying precision-based therapies^4, 12^.

Recent genomic studies in osteosarcoma have suggested that metastatic clones do not correspond to the dominant clones present in the primary tumor,^13^ and osteosarcoma may evolve via parallel evolution,^14^ with evidence for both monoclonal^15^ and polyclonal synchronous seeding of metastases^13, 14^. Copy number alterations in consensus driver genes, such as *MYC* and *CDK4* were found to be likely early events^15^. Cisplatin-induced mutagenesis has also been highlighted as a potential driver of treatment resistance in recurrent osteosarcoma^15^. Despite these initial insights into the clonal heterogeneity of osteosarcoma, the extent to which neoadjuvant chemotherapy affects clonal selection in patients with a poor response to chemotherapy and the degree to which copy number alterations evolve from diagnosis to relapse remains unclear.

To address these open questions about clonal selection and heterogeneity in osteosarcoma, we performed whole-genome sequencing of 37 spatially and temporally separated tumor samples from eight patients with osteosarcoma who had a poor response to neoadjuvant chemotherapy (<90% necrosis). We describe spatial intermetastatic heterogeneity and temporal clonal evolutionary processes, with a focus on identifying and tracking unique copy number clones from diagnosis through relapse.

## Results

### Analysis of Single Nucleotide Variants reveal limited driver gene heterogeneity in temporally and spatially distinct osteosarcoma sample

To analyze clonal evolution and intratumoral heterogeneity in osteosarcoma, we performed whole-genome sequencing (WGS) of tumor tissues from multiple spatially and temporally distinct samples from eight individuals with relapsed/refractory osteosarcoma. DNA was extracted from 84 samples collected from 10 patients. After initial quality control, we sequenced 62 unique tumor samples from eight patients with WGS to a target depth of 80x (Supplementary Fig. 1). Of these eight patients, four had localized disease at diagnosis (OSCE4, OSCE5, OSCE6, OSCE9) and four had metastatic disease at diagnosis (OSCE1, OSCE2, OSCE3, OSCE10) (Fig. 1A), and the age at diagnosis was 11-27 years (four girls and four boys, Fig. 1C). All patients were treated at the Memorial Sloan Kettering Cancer Center and received methotrexate, cisplatin, and doxorubicin (MAP) chemotherapy (subsequent post-procedure treatment, Fig. 1A). Seven of the eight patients had a poor response to neoadjuvant chemotherapy (<90% necrosis at the time of primary resection, Fig. 1D), while OSCE5 had an upfront resection; therefore, the response to therapy could not be evaluated.

**Figure 1.**
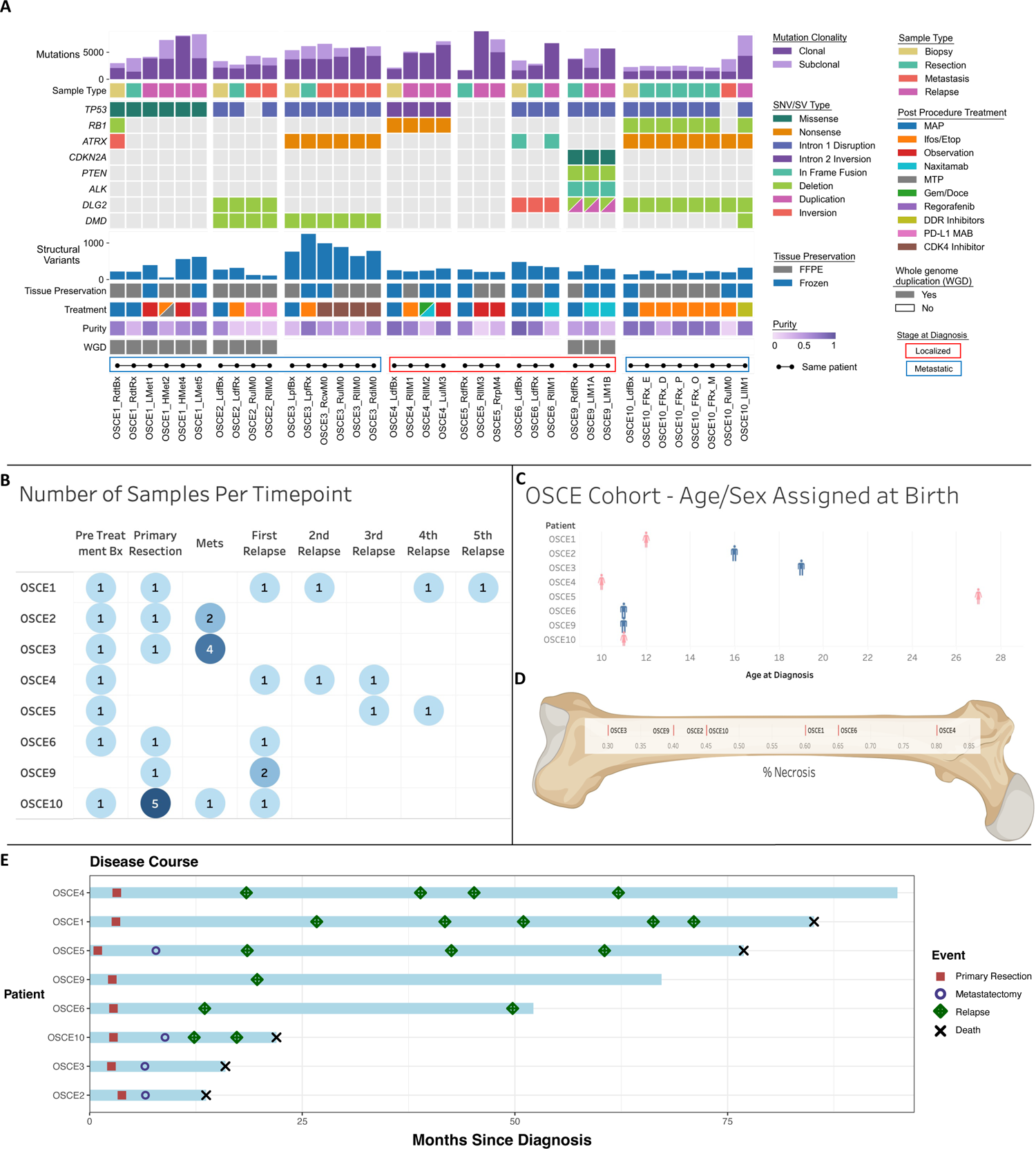
Characteristics of the patients and samples included in the analysis cohort. **A,** Oncoprint of sample and patient level details for each patient. Samples from the same patient are connected by dots and lines on bottom of figure and are in chronological order of time obtained (earliest on left). Sample Name Key: Bx = biopsy, Rx = Resection, Sample Ending in 0 = metastatic site present at diagnosis, sample ending in a number >0 indicates number of relapses. L/R = laterality, d = distal, p=proximal, f=femur, t=tibia, cw=chest wall, H=heart, ul=upper lobe, ll=lower lobe, di=diaphragm, rp=retroperitoneal, l=lobe. **B,** Summary of the number of samples per timepoint for each patient, darker shades of blue represent higher number of samples at a respective time point. **C,** Age and sex assigned at birth for each patient, patient is on the x-axis, age is on the y-axis, and sex assigned at birth is plotted on the chart as blue for male and pink for female. **D,** Percent necrosis at time of primary resection for each patient (Note OSCE2/OSCE10 both have 45% necrosis). **E,** Patients are listed on the y-axis and are ordered from longest disease course at the top to shortest at the bottom of the figure. The light blue bars represent length of disease course in months. Events are marked as depicted in the legend with different shapes and colors and plotted along the disease course bar at the time in months that the event occurs.

After sequencing was completed, we reviewed the quality of the sequencing data (purity, sequencing coverage) and determined that 37 of the 62 (59%) samples, similar to other studies^16^, met our quality control requirements to proceed with further downstream analysis (Supplementary Fig. 1). Of these 37 samples, 17 came from primary sites, with seven of eight patients having a pretreatment sample, six of which were biopsies and one (OSCE5), which was a pretreatment primary resection (Fig. 1A/B). Of the seven patients who did not undergo an upfront resection, all had at least one on-therapy resection sample, and one patient (OSCE10) had multi-region sampling from the primary tumor (Fig. 1A/B). The other 20 samples came from metastatic sites, 15 of which were from lung metastases, and 7 of the 20 were metastatic sites that were present at diagnosis (Fig. 1A/B). Fresh frozen samples accounted for 15 of the 37 samples selected for downstream analysis, while the remaining 22 were formalin fixed paraffin embedded (FFPE) samples (Fig. 1A). The median purity of fresh frozen samples was 0.75 compared to 0.46 of FFPE samples (Fig. 1A). All eight patients in this analysis had matched normal blood samples sequenced at a target depth of 40x.

The clinical courses of OSCE2, OSCE3, and OSCE10 were defined as refractory disease with progression on MAP (methotrexate, doxorubicin, cisplatin) chemotherapy and extremely virulent disease courses, with time from diagnosis to death of 1.08, 1.3, and 1.75 years respectively (Fig. 1E). OSCE1 and OSCE5 had long protracted relapsing and remitting disease courses, with a time from diagnosis to death of seven and six years, respectively (Fig. 1E). OSCE4 also had a relapsing and remitting disease course but is currently in remission eight years after diagnosis (Fig. 1E). OSCE6 is alive 4.5 years after diagnosis but has had a recent recurrence (Fig. 1E). OSCE9 is 4 years out from the initial metastatic recurrence (Fig. 1E).

After filtering and germline subtraction, whole-genome sequencing data identified between 1684 and 16,215 single nucleotide variants (SNVs) per sample (Fig. 1A). Of these, there were between 12 and 181 coding nonsynonymous SNVs per sample (median = 54, Supplementary Fig. 2A). The average number of nonsynonymous SNVs in primary site samples was 42, compared to 73 in metastatic or relapsed samples. SNVs were clustered across all samples for each patient using the DeCiFer^17^ algorithm, which determines the descendant cell fraction (DCF) of all SNVs for a given cluster in each patient (analogous to the cancer cell fraction but accounting for potential mutation losses^17^). Following previous approaches^17^, SNVs were categorized as clonal if they belonged to a cluster in a sample with a DCF ≥ 90%, and subclonal if they belonged to a cluster with a DCF < 90%. The proportion of clonal SNVs ranged from 35.7-100% across all samples (median = 68.2%), with relapse samples having the highest proportion (median = 92.7%) of clonal SNVs compared with biopsy, resection, and metastatic samples (median = 66.1%, 64.7%, and 63.6%, respectively) (Fig. 1A and Supplementary Fig. 1B). Likely functional driver gene SNVs^18^ were identified in five of the eight patients, including genes known to be frequently mutated in osteosarcoma, such as *TP53*, *ATRX*, *RB1*, and *CDKN2A* (Fig. 1A)^5–9^. These driver genes were clonal (shared) across all samples for each patient, and no new SNVs were likely functional drivers that were unique to any metastatic or recurrent samples (Fig. 1A, Supplementary Fig. 3A).

SNVs were clustered across all samples for each patient using the DeCiFer^17^ algorithm, which determines the descendant cell fraction (DCF) of all SNVs for a given cluster in each patient (analogous to the cancer cell fraction but accounting for potential mutation losses^17^). Following previous approaches^17^, SNVs were categorized as clonal if they belonged to a cluster in a sample with a DCF ≥ 90% and subclonal if they belonged to a cluster with a DCF < 90%. The proportion of clonal SNVs ranged from 35.7-100% across all samples (median = 68.2%), with relapse samples having the highest proportion (median = 92.7%) of clonal SNVs compared with biopsy, resection, and metastatic samples (median = 66.1%, 64.7%, and 63.6%, respectively) (Fig. 1A and Supplementary Fig. 1B).

Since there were no new SNVs in consensus driver genes unique to relapse or metastatic samples, we next examined structural variations and copy number alterations. Structural variants (SVs) in consensus driver genes were shared across all samples for each patient. *TP53* structural variants involving intron 1 were observed in 5/8 patients (OSCE2, OSCE3, OSCE6, OSCE9, and OSCE10; Fig. 1A). A *TP53* intron 2 inversion was observed in OSCE4 (Fig. 1A). There was a *TP53* SNV in OSCE1 and while there was no *TP53* SNV or SV found in OSCE5 (Fig. 1B), there was amplification of *MDM2*, an important negative regulator of *TP53*. In OSCE1, there was a deletion event in *RB1* and an inversion in *ATRX* in the pretreatment sample (RdtBx) that was not seen in the primary resection or relapse samples (Fig. 1A). Disruptions in *DLG2*, a bone tumor supressor gene^19^, were observed in 4 patients (OSCE2, OSCE6, OSCE9, and OSCE10; Fig. 1A). Deletion events in *DMD*, a gene that has been linked to aggressive behavior in human cancers, and is believed to have a potential role as a tumor suppressor, were observed in the three refractory cases (OSCE2, OSCE3, and OSCE9) and were present in all samples for OSCE2 and OSCE3; however, they were only detected in the relapse sample in OSCE10 (Fig. 1A). OSCE9 was found to have a deletion event in *PTEN* and an in-frame fusion event in *ALK* (Fig. 1A).

### Subclonal Somatic Copy Number Alterations Emerge and Dominate in Recurrent Osteosarcoma

A high prevalence of somatic copy number alterations (CNAs) in osteosarcoma has been reported previously^5–9^. Therefore, we used the HATCHet^20^ algorithm to infer both allele and clone-specific CNAs as well as their relative proportions across multiple samples from the same patient. The average tumor ploidy for each sample ranged from 1.7 in OSCE10 to 3.15 in OSCE3 (Supplementary Fig. 2B). HATCHet^20^ identified subclonal CNAs in all but one patient (OSCE2) and whole genome duplications present at diagnosis in three of the eight patients (OSCE1, OSCE2, and OSCE9; Fig. 1A). Among the seven patients with subclonal CNAs, six were identified as having two major copy number clones (Fig. 2A-E, Supplementary Fig. 4A), and one patient (OSCE10) had three distinct copy number clones (Fig. 2F). In four of the seven patients (OSCE1, OSCE4, OSCE6, and OSCE10), multiple distinct copy-number clones were identified to be simultaneously present in the primary tumor (Fig. 2A, 2B, 2D, 2F), but only one of these subclones emerged and dominated at subsequent relapses. In two of these patients (OSCE1, OSCE6), the emergence of this clone was observed in the primary resection sample when compared to the pretreatment biopsy for each patient (Fig. 2A, 2D). In OSCE5, a new copy number clone emerged in the late relapse samples, which was not identified as being present in the pretreatment sample (Fig. 2C). However, when combining the analysis of mutations and CNAs (see Online Methods), the dominant SNV-based clone (which shared the dominant copy number profile of the late emerging copy number clone) in the relapse samples was found to be sub-clonal in the pretreatment sample, providing evidence that the dominant copy number clone in the relapse sample for OSCE5 was present at diagnosis (Supplementary Fig. 5A). In OSCE10, no subclonal copy number aberrations were identified in the pretreatment sample; however, a subclonal copy number clone was detected in the primary resection sample that emerged and dominated at relapse in this patient (Fig. 2F). In summary, we found that in most cases, a minor subclone present in the primary tumor emerged and dominated in patients with relapsed disease.

**Figure 2.**
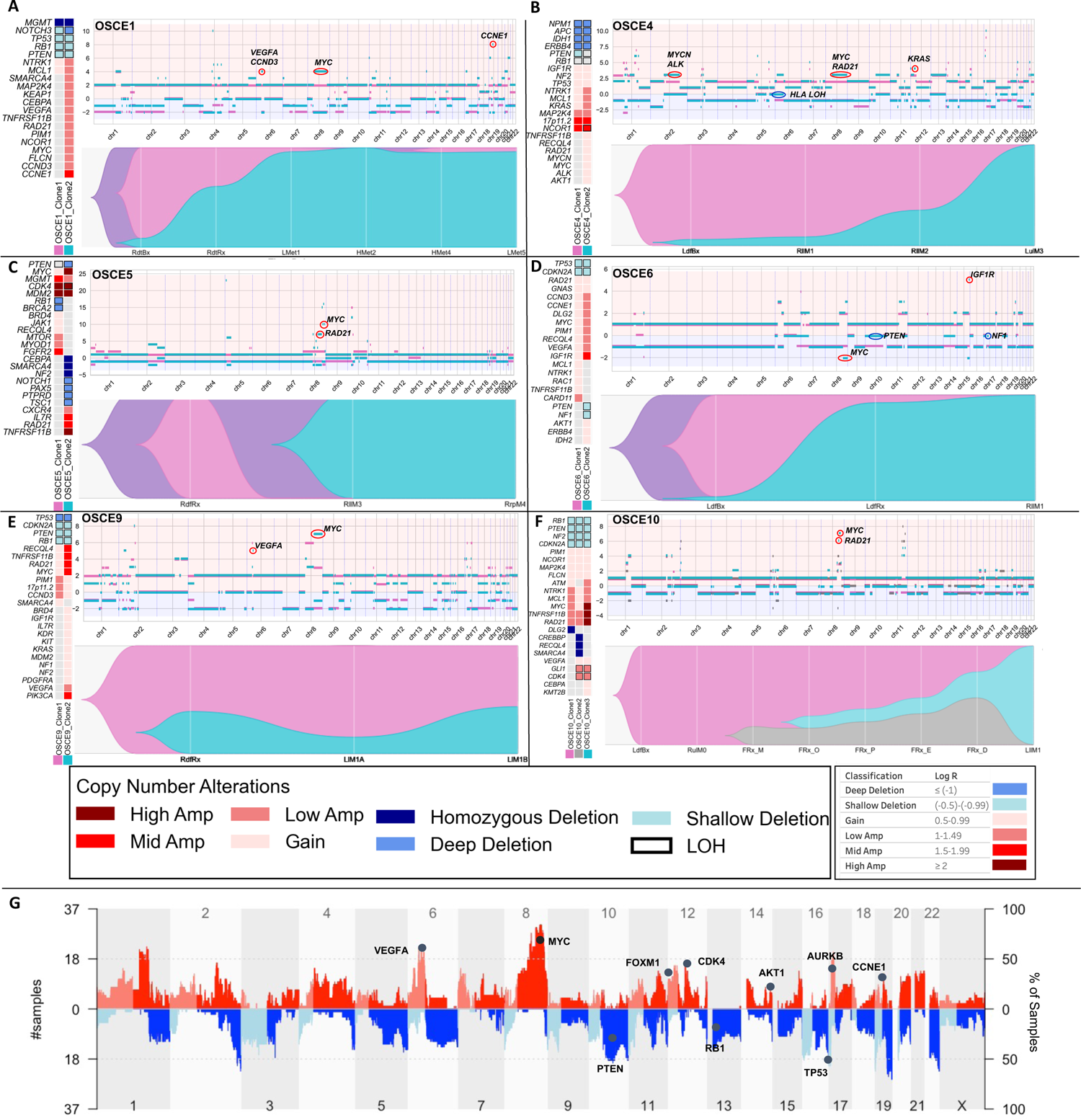
Subclonal copy number clones emerge at relapse. **A, B, C, D, E, and F,** For each patient there is a panel of three figures. The figure on the left is an oncoprint featuring clone specific copy number alterations in recurrently altered genes of interest in osteosarcoma. The top figure is a plot of allele specific copy number alterations for each clone with significant events for each clone circled and highlighted (note y-axis scales are unique for each patient). Clone 1 is the magenta clone, clone 2 is the teal clone, and clone 3 in OSCE10 is the gray clone. The major allele is plotted above 0 and the minor allele is plotted below 0. The bottom figure in each panel is a TimeScape plot of the prevalence of each clone at different timepoints throughout a patient’s disease course. **G,** Combined genome-wide copy number alterations across all patients in the cohort with recurrently altered genes highlighted.

In 2/3 refractory cases (OSCE2, OSCE3), there was no subclonal copy number clone that was identified in the primary that emerged and dominated in metastatic or relapse samples. In OSCE2, only a single major copy number clone was identified; in OSCE3, two copy number clones were identified in the pretreatment sample, with the copy number subclone emerging and dominating in one of the metastatic sites but in mixed proportion in the three other metastatic sites. OSCE9 did not have a pretreatment sample for comparison but did show two copy number clones in mixed proportions in the primary resection sample and the two relapse sites.

Determinations regarding branched vs. parallel evolution (depicted in the TimeScape plots in Fig. 2A-F, Supplementary Fig. 4A) for patients with ≥2 copy number clones were based on a review of loss of heterozygosity (LOH) events in the dominant metastatic or recurrent copy number clone, using the rationale and methods outlined by Watkins et al^21^. Branched evolution with the emergence of the treatment resistant copy number clone from the dominant pretreatment copy number clone (clone 2 emerging from clone 1) was observed in three patients (OSCE4, OSCE9, and OSCE10). Parallel evolution, where the pretreatment and treatment resistant copy number clones share the same parent clone (clone 1 and clone 2 share the same parent clone) but evolve in parallel, was seen in four patients (OSCE1, OSCE3, OSCE5, OSCE6). LOH events were common in tumor suppressor genes such as *TP53, RB1,* and *PTEN* and were mostly shared between clones for each patient, with a median of 77.95% of LOH events shared between all clones (range 18.9%-86.82%, Supplementary Fig. 4C).

### Copy number amplifications in recurrently altered oncogenes in osteosarcoma characterize chemoresistant copy number clones

Cohort-wide copy number alterations reflected what has been previously described in osteosarcoma^4^, with gains and amplifications seen in *VEGFA*, *MYC*, *FOXM1*, *CDK4*, *AKT1*, *AURKB*, and *CCNE1*, and loss/deletion events in *PTEN*, *RB1*, and *TP53* (Fig. 2G). In contrast to previously identified SNV drivers, the relative proportions of alterations across tumor cells changed with time. Clones were classified as chemoresistant if they were present in the primary site and became dominant at relapse, and chemosensitive if it was the dominant clone in the primary site and became subclonal or eliminated in metastatic or relapse sites. When comparing the genomic alterations between copy number clones for each patient, deletion or LOH events in tumor suppressor driver genes were often clonal in patients found to have ≥ 2 copy number clones (Fig. 2A-F, Supplementary Fig. 4A). In the 5 patients with clear emergence of a chemoresistant copy number clone (clone 2 in OSCE1, OSCE4, OSCE5, OSCE6 and clone 3 in OSCE10), the resistant clone had a higher degree of *MYC* gain or amplification then the dominant chemosensitive clone at diagnosis (Fig. 2A-F, Supplementary Fig. 5B). In OSCE10, high-level (log2 ≥ 2) MYC-amplified clone 3 emerged at the time of primary resection and dominated at relapse (Fig. 2F). Notably, this treatment-resistant clone was present in 4/5 multi-region samples from the primary resection, suggesting intratumoral heterogeneity regarding copy number alterations depending on the site sampled. In addition to *MYC*, amplification of *CCNE1*/*CCND3* (OSCE1, OSCE6; Fig. 2A/D), *KRAS* (OSCE2, OSCE4; Supplementary Fig. 4B, Fig. 2B), *IGF1R* (OSCE6, Fig. 2D), *CDK4* (OSCE10, Fig. 2F), *VEGFA* (OSCE1, OSCE6, OSCE9; Fig. 2A/D/E), and LOH at HLA (OSCE4, Fig. 2B and Supplementary Fig. 4D) uniquely characterized treatment-resistant or metastatic copy number clones in this cohort.

### Subclonal selection/emergence evident at the time of primary resection

Three patients within our cohort had a pretreatment sample on therapy resection, and at least one relapse sample (OSCE1, OSCE6, OSCE10), which provided an opportunity to analyze the effect of neoadjuvant chemotherapy on subclonal copy number selection at the time of primary resection and whether this selection is reflective of the dominant clone at the time of relapse. OSCE1 and OSCE6 share a similar pattern, where a copy number subclone present at diagnosis emerges as the dominant clone at the time of primary resection and continues to dominate at the first and subsequent relapses. Chemoresistant clone 2 was characterized by mid-level (log2=1.5-1.99) *CCNE1* amplification in OSCE1 (Fig. 2A) and mid-level *IGF1R* amplification in OSCE6 (Fig. 2D). A slightly different pattern emerged in the refractory case of OSCE10, in which only a single copy number clone was present at diagnosis. The primary resection sample underwent multi-region sequencing of five spatially separated sites, which revealed the emergence of two new copy number clones, with clone 2 present in all five samples and clone 3 in 4/5 samples. Clone 3 then became the dominant clone in the first relapse specimen, characterized by high-level *MYC* amplification.

### Timing of Copy Number Gains Reveals Large Bursts of Copy Number Gains Before Diagnosis

The HATCHet algorithm also infers whole-genome duplication (WGD) events jointly across all samples for each patient. Of the five patients with average ploidy ≥ 2.5 (OSCE1, OSCE2, OSCE3, OSCE4, OSCE9), WGD was identified in OSCE1, OSCE2, and OSCE9 (Fig. 1A and Supplementary Fig. 2B). The timing of these WGD events as well as other chromosomal duplications can be determined in “molecular time” using previously described methods^23, 24^ which compares the number of duplicated vs non-duplicated mutations to estimate the timing of each duplication (Supplementary Fig. 6). For the patients who were determined to have WGD events, these appeared to be late events for each respective patient (median point mutation time (pmt) range = 62-79%, Fig. 3F), compared to the two patients with ploidy ≥2.5 without WGD (patients with ploidy < 2.5 did not have enough duplication or LOH events to analyze), which appeared to have early synchronous duplications (median pmt range = 16-25%, Fig. 3F). The two patients with the lowest ploidy (OSCE10 and OSCE6) also had the smallest primary tumors (Fig. 3E), whereas all large primary tumors (≥300 cm^3^) had ploidy ≥ 2.5 (OSCE2, OSCE9, OSCE3). For the five patients with ploidy ≥ 2.5, we analyzed their earliest primary site samples to assess the natural history of these duplication events in the context of tumorigenesis.

**Figure 3.**
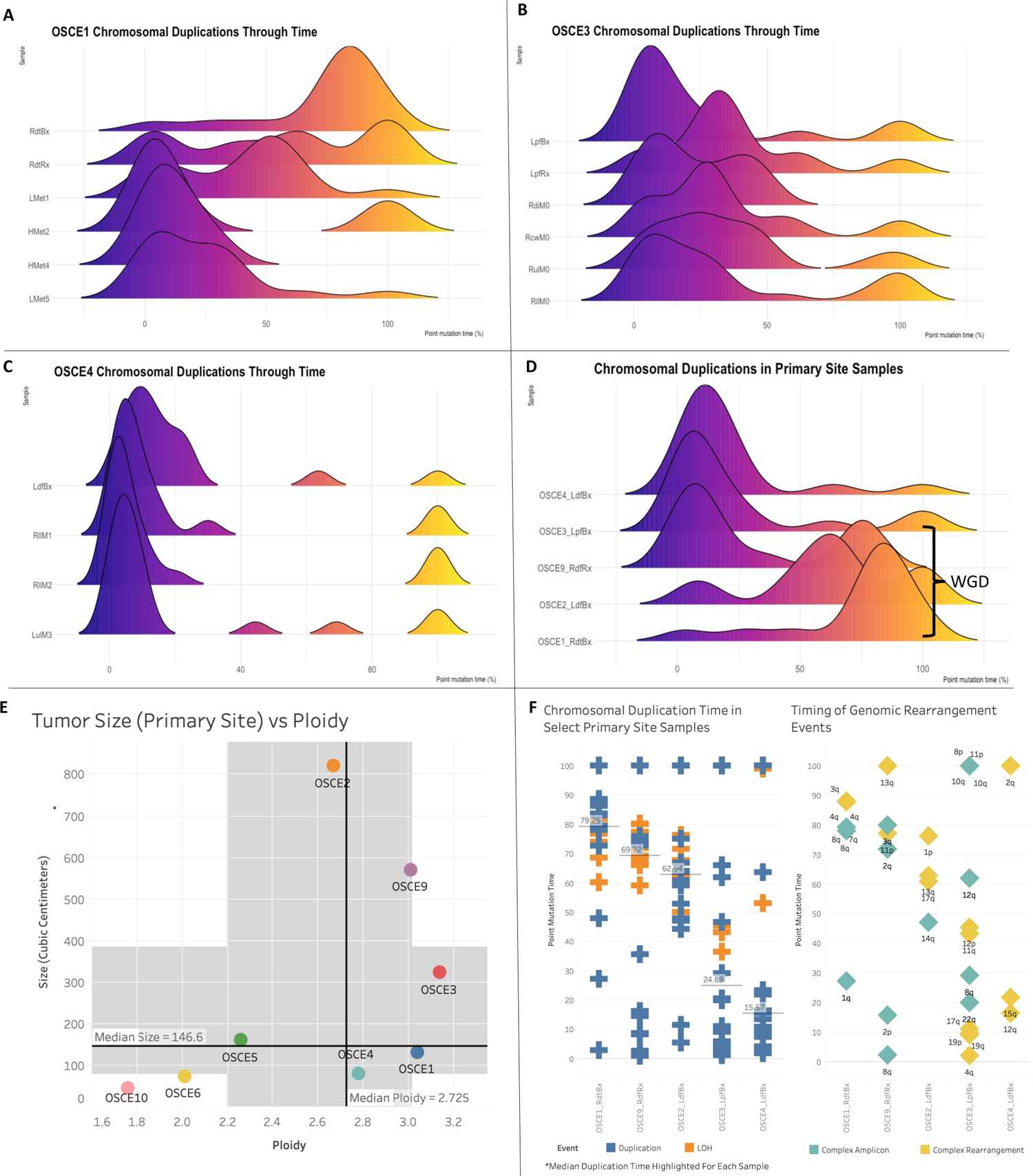
Chromosomal duplication timing analysis reconstructs evolutionary past of genomic instability events. **A, B, C,** Ridgeline plots of the density of duplication events over molecular time for each sample for the selected patients. Notably the highest peak in duplications occurs before diagnosis. **D,** Chromosomal duplication timing of 5 primary site samples. Whole genome duplication events as called by HATCHet appear to be a late event in our cohort. **E,** Plot of tumor size (by volume) verse ploidy called by HATCHet. Median size (y-axis) and ploidy (x-axis) values are plotted with dark black lines, with the shaded gray areas representing the range between the lower and upper quartile for each metric. **F,** Plot of duplication and rearrangement events in molecular time. Left side of figure is plot of duplication events in blue and LOH events in orange for select primary site samples. The median duplication time is highlighted for each sample. The plot on the right side of the figure are complex amplicon events in teal and complex rearrangement events in yellow. Y-axis for both plots is molecular time. The samples are in the same order for each plot. Each plotted event represents an affected chromosomal arm.

Most of these duplication/LOH events clustered in a single burst of events (Fig. 3D) and were associated with complex rearrangement (chromoplexy, chromothripsis) or complex amplicon^25^ (Tyfonas, breakage fusion bridge, double minute) events (Fig. 3F). When comparing longitudinal samples from the same patient (Fig. 3A-C), there was no subsequent burst that was greater than the primary site burst. Across the cohort, duplication events appeared to be fixed during tumorigenesis and had decreasing average molecular times when comparing biopsy/resection samples with metastatic/relapse samples (Supplementary Fig. 7A/C). LOH events were consistently “late” events, with a median molecular time of 82.16 across the cohort (compared to 21.72 for duplication events, Supplementary Fig. 7B/D).

### Heterogenous seeding patterns observed in metastatic and recurrent osteosarcoma

Clone-based phylogenies were created to explore the clonal architecture and track the spatial and temporal patterns of evolution. The SNVs across all samples for each patient were clustered using the DeCiFer^17^ algorithm. After clustering, each SNV was assigned to a clone with an estimated descendant cell fraction (DCF) per sample, and clone-based phylogenetic trees were then constructed using the CALDER^26^ algorithm, allowing for the assessment of modes of metastatic seeding and dissemination. The median number of clones per patient was eight (range = 5-12, Supplementary Fig. 8I). At the patient level, there was a heterogeneous mix of dissemination patterns (Fig. 4A-D, Fig. 5A-D, and Supplementary Fig. 8A-H), with complete agreement between clone and sample-based phylogenies (Supplementary Fig. 3A) regarding whether metastatic dissemination was monophyletic (all metastatic clones shared a common ancestral clone: Fig. 4A, 4C, 5E; Supplementary Fig. 6A, 6C, 6D, 6G, 6H)) vs. polyphyletic in origin (where ≥2 clones are present from distinct branches of phylogeny whose common ancestor represents the trunk of the primary tumor tree: Fig. 4B, 4D, 5B; Supplementary Fig. 6B, 6D, 6F). Of note, OSCE6 was the only patient among those with a monophyletic origin that had monoclonal seeding (one clone present in the sample, Fig. 4C and Supplementary Fig. 6H). The seeding and dissemination patterns were also assessed for each metastatic sample (Fig. 5E). In two patients (OSCE3 and OSCE5), there were examples of different modes of metastatic dissemination among spatially separated metastases from the same resection. In OSCE3 for example, extra-pulmonary metastases (RcwM0, RdiM0) demonstrate polyclonal (≥2 clones present in the sample) monophyletic patterns of dissemination, while the pulmonary metastases (RllM0, RulM0) demonstrate monoclonal monophyletic dissemination, with both samples sharing the same ancestral clone (Fig. 5C, Supplementary Fig. 6B, 6I).

**Figure 4.**
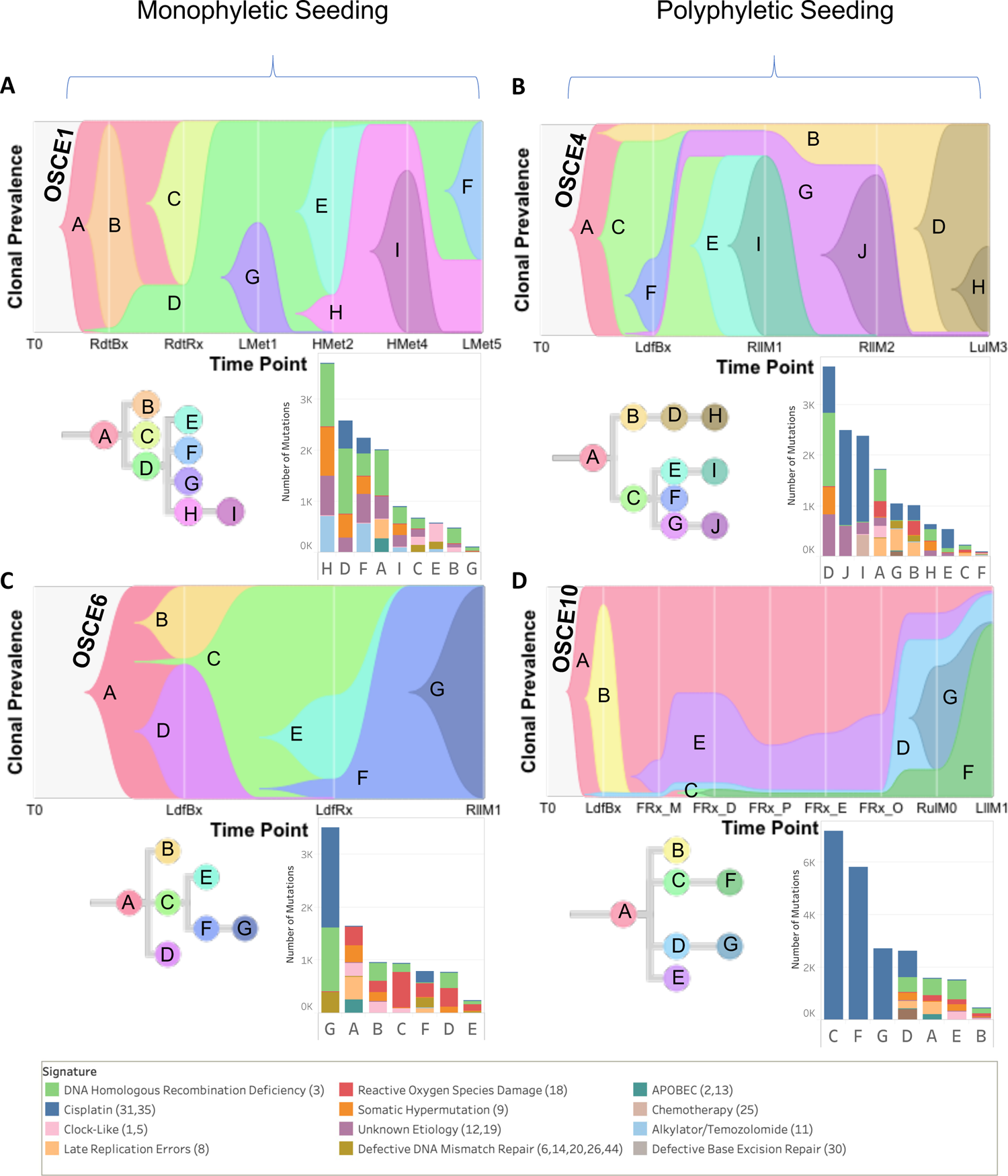
SNV based phylogenies highlighting temporal evolution with clonal mutational signature composition. **A, B, C, and D,** Upper figure in each panel is a TimeScape plot of the inferred evolutionary phylogeny, highlighting clonal proportions over time. The prevalence of different clones is shown over time on the vertical axis, with the different clones represented by different colors. The horizontal axis represents the timepoints, which are represented by gray lines. The evolutionary relationships between the clones are shown on the phylogenetic tree and in the TimeScape layout. The bottom right of each panel is a stacked bar plot of the total number of mutations assigned to each clone. Colors represent total number of mutations attributed to each mutational signature with color legend at bottom of figure. Patient level metastatic seeding patterns are denoted by the brackets at the top of the page.

**Figure 5.**
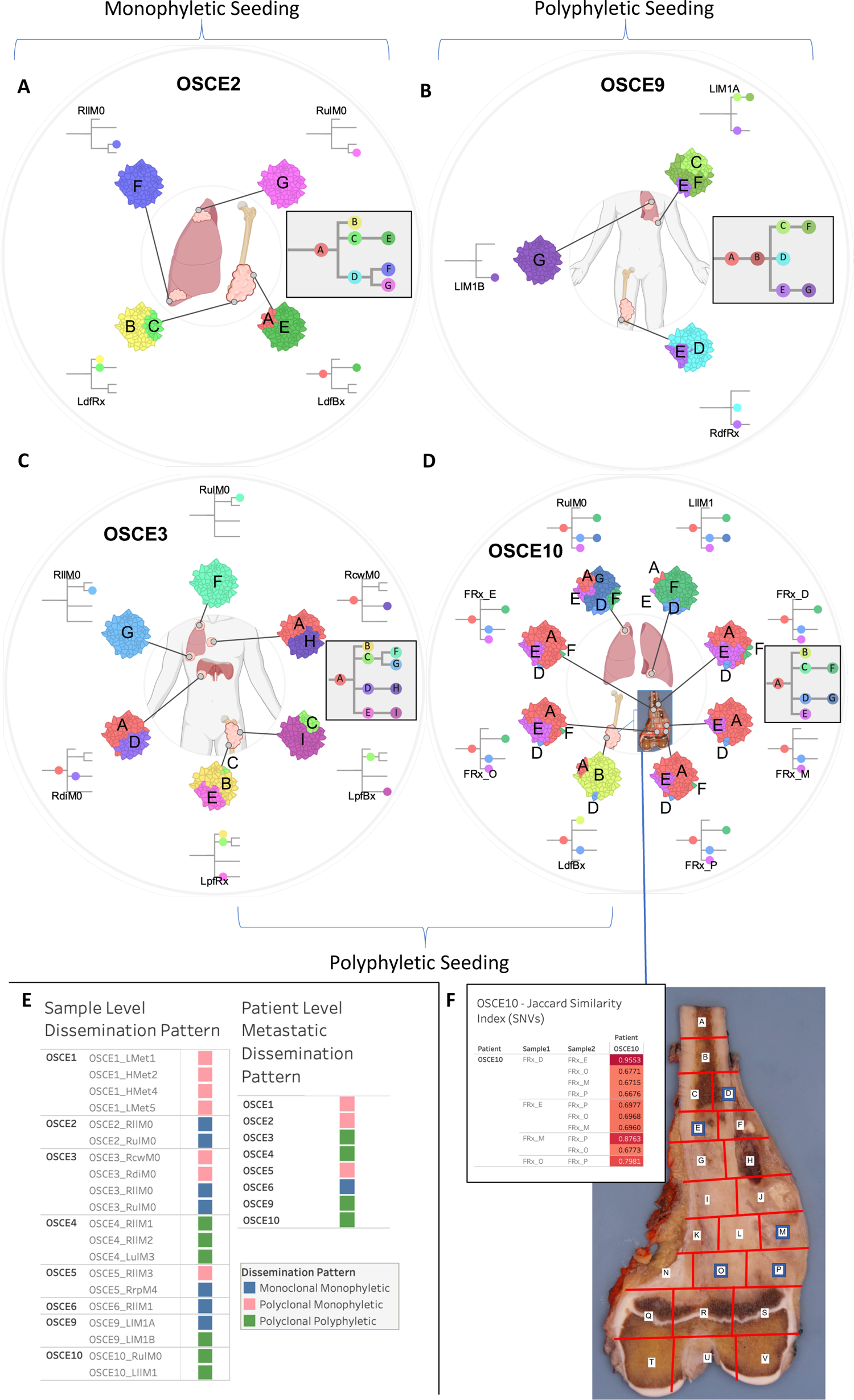
SNV based phylogenies highlighting spatial evolution and descriptions of metastatic seeding patterns at the sample and patient level. **A, B, C, and D,** Spatially and in some cases temporally distinct samples are indicated on the anatomic sites from where the sample originated. The colors represent different clones, and the phylogenetic trees show the evolutionary relationships between these clones. The prevalence of each clone at a particular site is proportional to the colored area of the cellular aggregate representation. **E,** Sample and patient level dissemination patterns are characterized in these charts. *Monoclonal dissemination*: single subclone within the primary tumor seeds one or more metastatic lesions, *polyclonal dissemination*: multiple distinct subclones from the primary tumor seed one or more metastatic lesions, *monophyletic origin*: all metastatic clones are derived from a recent common ancestor, *polyphyletic origin*: metastasizing clones are more similar to other subclones within the primary tumor than they are to each other. These descriptions can be considered at the sample level, focused on the clonal make up of a single metastatic site compared to the primary tumor, or taken as a whole, evaluating all spatially or temporally separated samples and how they relate back to the primary tumor. **F,** Multi-region sequencing was performed on a primary resection sample from OSCE10. Regions D, E, M, O, P were sequenced from the specimen grid depicted. A table of Jaccard similarity indexes based on shared SNVs for these samples is shown in the upper left inset.

**Figure 6.**
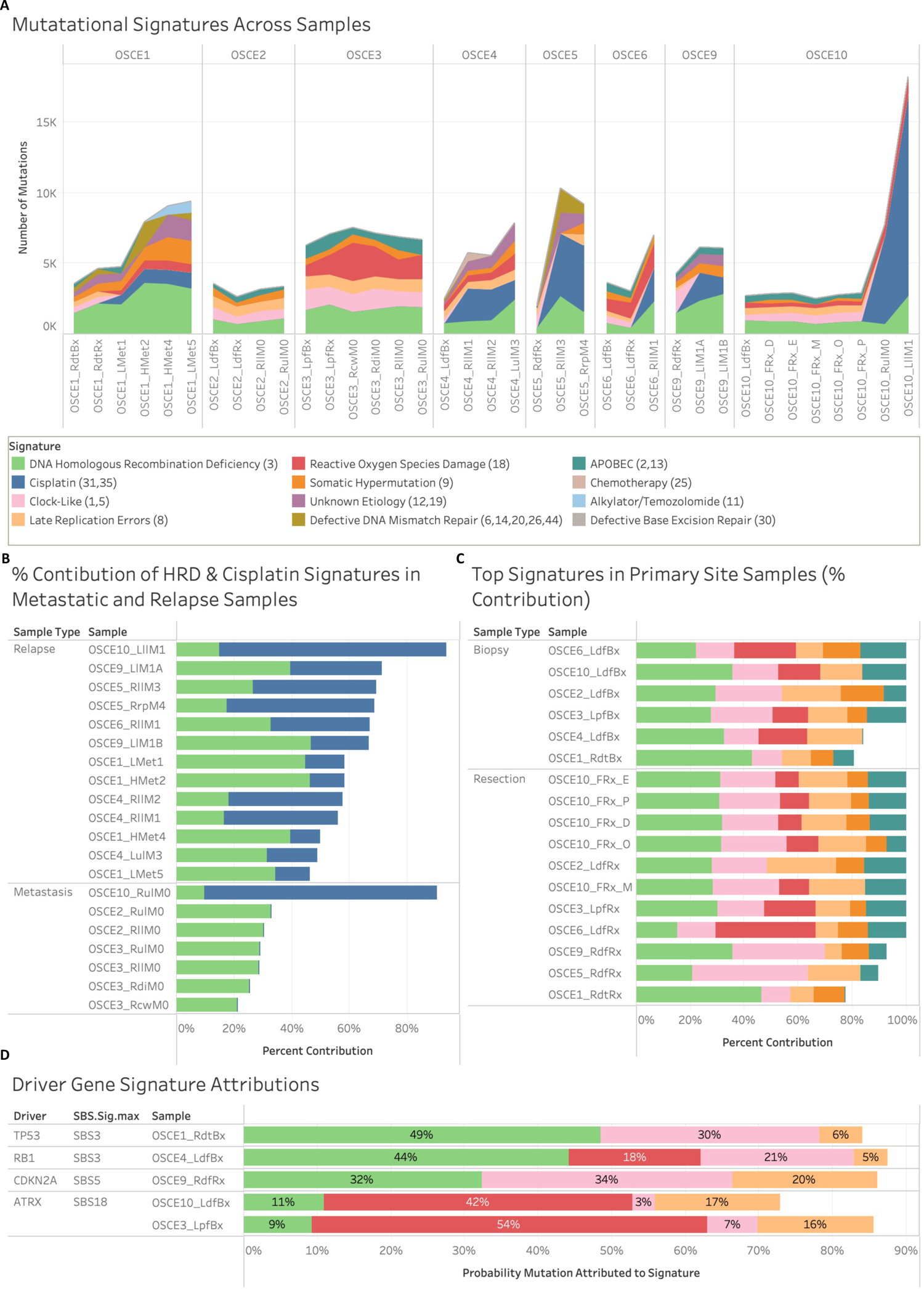
Mutational signature patterns across the cohort. **A,** Stacked line chart of mutational signature contribution by total number of mutations attributed to each signature. Colors represent the different signatures. **B,** Stacked bar chart of the relative contribution of HRD-related SBS3 (green) & cisplatin (blue) signature in metastatic and relapse samples. **C,** Stacked bar chart of the relative contribution of HRD (green), clock-like (pink), reactive oxygen species damage (red), late replication errors (beige), somatic hypermutation (orange), and APOBEC (teal), in primary site samples. **D,** Stacked bar chart of the probability that driver gene SNVs were attributed to a mutational signature. Pretreatment samples from patients with driver SNVs were included and the primary resection sample from OSCE9 since no pretreatment sample was available. Colors represent the different signatures.

### Limited Heterogeneity after Induction Chemotherapy

For OSCE10, we obtained a section of the primary tumor sample for multi-region sequencing after the patient had received 10 weeks of induction MAP chemotherapy. The specimen was mapped (Fig. 5F), DNA was extracted from the sections with viable tumor, and then five sections (E, D, M, O, P) with the highest DNA quantity/quality metrics underwent WGS. When examining the clonal architecture of each section (Fig. 4D/5F), each section had the same four clones (A, D, E, F), with truncal clone A dominating, except for FRx_M, which did not have clone F, which later became the clone that dominated at relapse. This pattern was also seen with the copy number clones, where clone 3 was present in all sections except for FRx_M and then dominated at relapse (Fig. 2F). When comparing Jaccard similarity coefficients based on SNV composition across the different sections, adjoining sections shared the highest coefficients (D/E=0.96, O/P=0.79, M/P=0.88, Fig. 5F), whereas non-adjoining sections had similar coefficients, regardless of distance from each other (range = 0.67-0.69, Fig. 5F). As highlighted previously, a single-sample sequencing strategy that sampled from FRx_M would have missed detecting the metastatic subclone that was present in the other four sections.

### Most new SNVs in relapsed disease are attributed to HRD-related SBS3 and cisplatin mutational signatures

To further characterize the potential drivers of clonal evolution, mutational signature analysis was performed for each clone (Fig. 4A-D, Supplementary Fig. 6A-D). In three patients, the largest clone by number of SNVs (OSCE5 – clone C, OSCE6 – clone G, and OSCE10 – clone C) had over half of the SNVs attributed to the DNA-damaging effects of cisplatin chemotherapy (Supplementary Fig. 9A). In four patients, the largest clone by number of SNVs (OSCE1 – clone H, OSCE3 – clone A, OSCE4 – clone D, and OSCE9 – clone G) had a plurality of SNVs in each clone attributed to single base substitution (SBS) 3, a genomic signature that has been associated with homologous repair deficiency (HRD, Supplementary Fig. 9A). Tumors with this signature are thought to have a BRCAness phenotype and exhibit features similar to those cancers with germline BRCA1 or BRCA2 mutations, even though no mutations in those genes have been identified. In OSCE2 and OSCE3, both refractory cases, the largest number of SNVs was assigned to the truncal clone, clone A, which had a high number of SNVs (OSCE2 – 1731/2234, OSCE3 – 2464/4022) associated with HRD-related SBS3 and late replication errors (Sig. 8). In patients with patient-level monophyletic seeding of metastases (OSCE1, OSCE2, OSCE5, OSCE6), where a single ancestral metastatic clone could be identified (OSCE1–clone D, OSCE2–clone D, OSCE5–clone C, OSCE6–clone C), there were no clear patterns regarding the signature composition identified (Supplementary Fig. 9B). OSCE1–clone D and OSCE2-clone D had a plurality of SNVs attributed to HRD-related SBS3, whereas OSCE5-clone C had the majority of SNVs attributed to cisplatin, and OSCE6 had the majority of its SNVs attributed to reactive oxygen species damage (Sig. 18). When looking at doublet base substitution (DBS) signatures at the clonal level, in clones with ≥10 DBS SNVs, cisplatin-associated DBS 5 was the largest contributor of DBS SNVs, accounting for 50% or more of the total SNVs in 17/26 clones (Supplementary Fig. 9C).

### Emergence of cisplatin and alkylator signatures helps time the formation of metastases

A simple method for ascertaining whether a given metastasis arose before or after treatment with cisplatin or an alkylator is to find clonal SNVs attributed to the respective signature in a metastatic tumor sample^27^. When looking at the dominant clone in the first relapse sample for patients with recurrent disease (OSCE1-Clone D, OSCE4-Clone E/I, OSCE5-Clone C, OSCE6-Clone C/F/G, OSCE9-Clone C/G, OSCE10-Clone C/F), there was a cisplatin signature present in each of these respective clones/clades, indicating that the metastases arose after therapy (Fig. 4A-D, Supplementary Fig. 6C, 6D). Additionally, in OSCE1, after the patient received chemotherapy with ifosfamide and etoposide, OSCE1-Clone H and OSCE1-Clone F had mutations attributed to the alkylator signature, SBS11 (Fig. 4A). In the three patients with refractory disease (OSCE2, OSCE3, and OSCE10), we can demonstrate which metastatic samples were present at diagnosis and which developed while on therapy. Within OSCE2, Clone F and Clone G were the dominant subclones in RllM0 and RulM0 respectively in OSCE2 (Fig. 5A, Supplementary Fig. 6A). Clone F had 181/614 mutations attributed to cisplatin and Clone G had no mutations attributed to cisplatin, evidence Clone F was seeded on therapy as opposed to prior to therapy. In OSCE3, there was no evidence of a cisplatin signature in any of the subclones (Supplementary Fig. 6B), indicating that all metastatic sites had developed prior to initiating therapy. In OSCE10, the metastatic sample (RulM0), which has polyphyletic and polyclonal seeding (Fig. 4D, 5D), showed that the dominant clone/subclone pair of D/G likely seeded on therapy, given that there is a cisplatin signature attributed to approximately half of the mutations (1027/2644) and in all the mutations in subclone G (2727/2727).

### HRD-related SBS3 and Reactive Oxygen Species Damage Linked to Driver Gene Mutagenesis

To identify the mutational processes most likely to be the origin of truncal driver gene SNVs, we calculated the likelihood that each individual SNV was caused by each signature, considering the mutation category and proportion of each mutational signature in the tumor genome^24^. To minimize the effect of treatment-related mutagenesis, we limited this analysis to the earliest primary site sample available for patients with truncal driver SNVs. Similar to the observations across all samples, HRD-related SBS3 had the highest probability of attribution in the *TP53* driven OSCE1 sample and the *RB1* driven OSCE4 samples (OSCE1 probability=0.49, OSCE4 probability=0.44, Fig. 6D), and a slightly lower attribution probability than the clock-like signature in OSCE9 (HRD-related SBS3 probability=0.32, clock-like probability=0.34, Fig. 6D). Both OSCE3 and OSCE10 had truncal *ATRX* SNVs, with reactive oxygen series damage accounting for the highest probability of attribution in both samples (OSCE3 probability=0.54, OSCE10 probability=0.42, Fig. 6D).

### Cisplatin associated hypermutation in a case of refractory osteosarcoma

When evaluating mutational signatures at the sample level, the two most prevalent SBS signatures across all samples were HRD-related SBS3 and cisplatin (Fig. 6A). In the 13 relapse samples, HRD-related SBS3 and cisplatin accounted for more than half of all the SNVs (Fig. 6B). In the 17 primary site samples, the HRD-related SBS3, clock-like (Sig. 5), late replications errors (Sig. 8), ROS damage (Sig. 18), and APOBEC (Sig. 2,13) accounted for 75%–100% of all mutations (Fig. 6C). In 6/7 metastatic samples, HRD-related SBS3, clock-like, late replication errors, somatic hypermutation, ROS damage, and APOBEC accounted for all SNVs (Supplementary Fig. 10A). In the metastatic sample thought to have emerged on therapy for OSCE10 (RulM0) and the subsequent relapse sample (LllM1), SNVs attributed to cisplatin accounted for 6226/7724 (86%) and 14276/18216 (78%) SNVs, respectively (Fig. 10A). In the six patients with relapsed disease, the number of mutations attributed to HRD-related SBS3 consistently increased from diagnosis to subsequent relapses, with at least 1000 new SNVs attributed to HRD-related SBS3 when comparing diagnostic and relapse samples (Supplementary Fig. 10C).

Doublet base substitution (DBS) signatures were also evaluated across the cohort (Supplementary Fig. 10B). DBS signature 5, which is associated with cisplatin, was detected in 21 samples; however, to filter out false positives, a threshold of ≥ 5 DBS signature 5 SNVs was used to confirm the absence of the signature. Using this filter, DBS signature 5 was present in 15 samples, all metastatic or relapse samples, with complete overlap with the 14 samples that had SBS cisplatin signatures 31 or 35. Only OSCE3_RdiM0 had a DBS cisplatin signature but not an SBS cisplatin signature (Supplementary Fig. 10D).

## Discussion

Tumor evolution and clonal heterogeneity have been increasingly recognized as major causes of therapeutic resistance to current anti-neoplastic therapies^28^. These findings extend our understanding of therapeutic resistance in spatially and temporally separated tumor samples from 8 patients with recurrent or refractory osteosarcoma. We found that while clonal driver gene SNVs and structural variants remain largely unchanged over the course of tumor progression, subclonal tumor populations with unique driver gene amplifications are present at diagnosis, emerge after treatment, and persist as the major clone at subsequent relapses.

Somatic copy number alterations are now increasingly recognized for their prognostic value over SNVs in multiple cancer types^29^. Oncogenic copy number alterations, while heterogeneous across osteosarcoma, represent potential therapeutic targets, given the lack of recurrent targetable SNVs or structural variants^4–8^. Our study revealed that in our four patients who had relapsed osteosarcoma with a matched pretreatment sample (OSCE1, OSCE4, OSCE5, OSCE6), there was a subclonal treatment resistant copy number clone that emerged as the dominant clone in the relapsed setting. Furthermore, in the two patients with both a pre-treatment sample and on-therapy primary resection (OSCE1, OSCE6), this treatment resistant clone clonally expanded after 10 weeks of neoadjuvant chemotherapy. We believe this finding has important implications for molecular profiling strategies in osteosarcoma, as it suggests that the primary resection sample, and not the pretreatment biopsy, is more reflective of the metastatic potential for a tumor than the pretreatment biopsy, due to the selection pressure of neoadjuvant chemotherapy. Achieving a cure in osteosarcoma requires the extinction of all cancer cells with a successful “first-strike’ strategy with maximum tolerated doses of cisplatin, doxorubicin, and methotrexate^30^. For patients for whom this first strike fails (poor necrosis at the time of primary resection), characterizing and targeting the treatment-resistant population of cancer cells using a second-strike strategy may prove to be an effective treatment strategy^30^. Our work highlights that molecular profiling of primary resection samples could allow for a more precise “genome-informed” approach^4^, aimed at targeting resistant copy number alterations, to augment MAP chemotherapy.

The emergence of subclonal copy number alterations in primary tumors to fully clonal alterations in metastatic or recurrent samples, as demonstrated in our study in 6/7 patients with recurrent/refractory disease and pretreatment samples, has been previously described in a subset of adult cancers where analysis of matched primary and metastasis was performed^21^. We found *MYC* gain/amplification to be enriched in the treatment-resistant clone in 6/7 patients with more than one clone. Previous studies have shown that *MYC* amplification is often enriched in metastatic sites^21^ and has been previously associated with poor outcomes and increased cell proliferation in osteosarcoma;^31–35^ however, a recent study questioned its prognostic significance^36^. Our study demonstrated that in patients with localized and metastatic disease at diagnosis, *MYC* amplification is subclonal in pretreatment samples and emerges after 10 weeks of neoadjuvant chemotherapy, highlighting the importance of sample timing when considering the prognostic value of *MYC* amplification. Furthermore, as we begin to define molecular risk categories within osteosarcoma, our work demonstrates that profiling of post-treatment primary resection samples may reveal previous subclonal amplifications in driver oncogenes; thus, this time point would be more informative when assessing metastatic potential. While multi-region profiling of post-treatment resection from OSCE10 revealed limited heterogeneity among the different sites, there was one site where the metastatic clone was not present, highlighting the potential risk of failing to profile the metastatic clone with single-sample strategies. When considering future sequencing approaches, pooling DNA/RNA extracts from multiple anatomically distinct tumor regions of the primary tumor could be a cost-effective way to improve DNA yield and variant detection, while providing a more complete picture of intratumoral heterogeneity^37^.

Our chromosomal duplication timing analysis revealed that gains for the same patient often clustered around the same time point, regardless of whether whole-genome duplication was present. These bursts of duplications occurred prior to diagnosis, and there were no comparable bursts of duplications in resection, metastatic, or recurrent samples that would reflect ongoing instability. In a pan-cancer cohort, synchronous bursts of copy number gain were found to occur in 57% of diploid samples and 78% of WGD samples^38^. We found that these clustered duplication events were associated with catastrophic complex genomic rearrangement and amplicon events that occurred before diagnosis, such as chromothripsis and tyfonas. In contrast to a previous multi-region osteosarcoma study^15^, we demonstrated that tumor ploidy remained consistent across all samples for each patient, which is likely because our copy number calls were inferred jointly across all samples for each patient, which can improve ploidy estimation in sample sets with wide ranges of purity^20^. These findings of pre-diagnostic duplication events followed by relative genomic stability support recent work that demonstrated that early catastrophic events are responsible for the structural complexity in the osteosarcoma genome, as opposed to sustained evolution and instability over time^39^. Whole-genome duplication was confirmed in a subset of patients and was found to be a late event in all three cases. Previous pan-cancer studies have found that WGD events are typically early events in a tumor’s molecular time history, but are often preceded by *TP53* inactivation^21, 38, 40^. Late WGD events have been described in a cohort of patients with hepatocellular carcinoma (HCC), and they are typically associated with larger tumors, leading to the conclusion that they may be the last step prior to rapid growth and expansion^24^. Our data support a macroevolutionary model of evolution in osteosarcoma^41^, with a large number of genomic aberrations acquired over a short period of time secondary to chromosomal instability events, followed by clonal selection, as opposed to ongoing evolution.

Large-scale genomic sequencing studies in osteosarcoma have revealed that there is significant inter-tumoral heterogeneity in osteosarcoma, with shared mutations typically in tumor suppressor genes rather than in targetable oncogenes^5–8^. We observed a heterogeneous mix of metastatic and recurrent seeding patterns in our cohort. We observed only one example of monoclonal, monophyletic dissemination in OSCE6, which is typically a result of a treatment-induced bottle-necking effect. There were three cases of polyclonal/monophyletic dissemination, in which multiple clones were present in the metastatic/recurrent samples, but they all shared a common ancestor, and there were four cases of polyclonal polyphyletic dissemination where multiple distinct clones from the primary tumor seeded a metastatic site. In previous studies, a *de novo* induced murine model of osteosarcoma demonstrated polyclonal seeding of metastases with ongoing parallel evolution^14^, while studies using longitudinal and spatially separated samples have yielded mixed results, demonstrating both polyclonal seeding with parallel evolution^13^ and monoclonal monophyletic seeding^15^ in a majority of the respective cases from each study. These studies were limited by the lack of pretreatment primary tumors; therefore, the analysis relied on comparing metastatic and recurrent samples to post-treated primaries in many cases.

We demonstrate that while cisplatin and HRD-related SBS3 are active mutagenic processes in osteosarcoma, accounting for most new mutations in relapsed disease, we found no new driver SNVs attributable to these signatures that could account for treatment resistance. The HRD-related SBS3 signature was conserved across all samples in our cohort, consistent with a recent pan-pediatric cancer study that found that 18/19 (95%) patients with osteosarcoma had mutations attributed to HRD-related SBS3^42^. The prevalence of the HRD-related SBS3 signature in osteosarcoma is comparable to BRCA1 deficient cancers, suggesting that drugs that target homologous recombination deficient cells, such as PARP inhibitors, may have therapeutic value in osteosarcoma, a concept currently being evaluated in a phase II clinical trial (NCT04417062)^43^. A commonly cited limitation of using signature-based assays to assess HRD is that they reflect the HRD state prior to sample acquisition and may not reflect the current state, where HRD may have been restored^44^. Our study demonstrated that in osteosarcoma, the number of mutations attributed to HRD-related SBS3 increases at each time point in patients with recurrent disease, suggesting that HRD continues to be an active mutagenic process after diagnosis.

The cisplatin signature was present in all relapse samples and metastatic sites that were thought to have developed during upfront therapy. The extent of cisplatin-induced mutagenesis has been previously described in osteosarcoma, where it was found that cisplatin therapy could potentially increase the mutational burden by two-fold^15^. Although we also found that cisplatin therapy led to large increases in mutational burden in recurrent samples, none of these mutations were likely drivers of treatment resistance, which is consistent with previous studies in patients with platinum-resistant ovarian cancer^45^ and osteosarcoma^15^. Although we cannot account for copy number alterations or structural variants induced by cisplatin, recent cell line work in cisplatin exposed esophageal and liver tumors, found few copy number alterations or structural variants, suggesting that cisplatin does not contribute significantly to genomic instability^46^.

Our findings highlight that the chemoresistant population of tumor cells in osteosarcoma is subclonal at diagnosis and is characterized by unique oncogenic amplifications. As our ability to target these oncogenic amplifications improves, future studies aimed at identifying these oncogenic drivers during upfront therapy may be an effective strategy to eliminate chemoresistant tumor cells and improve survival.

## Methods

### Patient consent and tissue processing

This study was approved by the Institutional Review Board of the Memorial Sloan Kettering Cancer Center (New York, NY, USA) and conducted in accordance with the Declaration of Helsinki. Informed written consent was obtained from each subject or guardian. Tumor samples and matched normal samples were collected from 10 patients with a pathologically confirmed diagnosis of osteosarcoma, who were identified both retrospectively and prospectively for those who had their tumor banked at diagnosis and at least one other time point. Only patients who consented to an IRB-approved blood and tumor collection protocol were eligible for tumor sequencing. Fresh tumor samples were procured from the operating room in a sterile container. The tissue was processed using scalpels and divided into pea-sized pieces before being stored at −80°C. Frozen tissue samples from several patients were also available through our Precision Pathology Biobanking Center and were acquired using the same protocol. Additionally, in several patients, archival tumor specimens in the form of formalin-fixed paraffin-embedded (FFPE) specimens were obtained from both the internal and external pathology departments using the same protocol. Only FFPE samples that were not subjected to harsh decalcification techniques were selected. FFPE samples that had been decalcified using EDTA were deemed appropriate for further downstream analysis.

Each frozen tissue sample was submitted to our pathology core, where it was embedded in Tissue-Tek optimum cutting temperature compound and sectioned at 5–10 mm on a Leica Cryostat to create a hematoxylin and eosin–stained (H&E) slide for review. Each FFPE sample was sectioned using a Leica Microtome. H&E slides were evaluated by a trained pathologist to determine tumor content. After pathologic review, tumor samples were isolated via a 21-gauge punch, curl biopsy, or macro-dissected from sectioned slides to remove non-neoplastic components. The neoplastic component of each tumor underwent genomic DNA extraction using a Qiagen DNAeasy Blood and Tissue Kit and protocol, whereas the FFPE samples were extracted using a QIAamp DNA FFPE Tissue Kit and protocol.

PBMCs utilized for matched normal sequencing were brought up to 15mL volume in cold PBS and isolated with the DNeasy Blood & Tissue Kit (QIAGEN catalog # 69504) according to the manufacturer’s protocol and incubated at 55°C for digestion. DNA was eluted in 0.5X Buffer AE.

### Whole-genome sequencing and alignment

DNA quantification, library preparation, and whole-genome sequencing were performed using the Integrated Genomics Operation at the Memorial Sloan Kettering Cancer Center (New York, NY). After PicoGreen quantification and quality control using an Agilent BioAnalyzer, 131-500ng of genomic DNA was sheared using an LE220-plus Focused-ultrasonicator (Covaris catalog # 500569), and sequencing libraries were prepared using the KAPA Hyper Prep Kit (Kapa Biosystems KK8504) with modifications. Briefly, libraries were subjected to a 0.5 × size selection using aMPure XP beads (Beckman Coulter catalog # A63882) after post-ligation cleanup. Libraries with < 500 ng of genomic DNA were amplified using 5-6 cycles of PCR and pooled equimolar amounts. Libraries containing at least 500 ng of genomic DNA were not amplified. Samples were run on a NovaSeq 6000 in a 150bp/150bp paired end run, using the NovaSeq 6000 SBS v1 Kit and an S1, S2, or S4 flow cell (Illumina). The average number of read pairs per normal was 614 million and the average number of read pairs per tumor was 1.3 billion.

### Whole Genome Sequencing Pipeline

Whole genome sequencing data were analyzed using the ISABL^47^ platform, with methods previously described in detail^48^. The additional downstream analyses are described below.

### Single Nucleotide Variant Filtering

For all eight patients, single nucleotide variants (SNV) were called in triplicate by MuTect^49^, Strelka^50^, and Caveman^51^. Only mutations that had a “PASS” flag, were called by at least two mutation callers and observed in less than 2% of the reads of the matched normal sample with 10x coverage were considered for further analysis. It is notoriously difficult to extract high quality DNA from the osteoid matrix that surrounds viable tumor cells, often leading to sequenced samples with low purity. Across the eight patients, a mix of samples was prepared as either fresh-frozen or FFPE. FFPE samples are thought to be inferior to fresh frozen samples, as the formalin fixation process results in nucleic acid fragmentation, DNA crosslinks, and deamination, leading to C>T mutation artifacts. As a result, downstream analysis of FFPE can be challenging when filtering out artifacts from a true positive. Aggressive filtering of low-allele frequency variants has been shown to increase SNV overlap when comparing matched FFPE and FF samples^52^. Given these assumptions, a custom filtering approach was utilized to maximize the ability to utilize low-purity FFPE samples within a mix of higher-quality FFPE samples and fresh frozen samples for each patient. For fresh frozen samples with an estimated purity of 20%, mutations with a variant allele frequency (VAF) less than 5 were filtered out. If a frozen sample had a purity of less than 20%, no additional filtering was applied, given the potential for filtering out subclonal mutations in an otherwise high-quality frozen tissue sample. For FFPE samples, we filtered out mutations with a VAF less than 20%. These thresholds for FFPE samples were then further purity adjusted, for example a sample with an estimated purity of 80%, would have mutations below a VAF of 0.16 (0.8 purity * 0.2 VAF filter) initially filtered out. Although this initial filtering for FFPE is strict, the advantage of having multiple samples per patient allows us to utilize the mutation calls in other high-quality samples for a patient to rescue mutations that may have been initially filtered in a low-quality sample. This is accomplished through a pile-up rescue, where all filtered mutations for a patient are combined and then specifically searched in the BAM file that was generated for each sample.

### Driver Gene Analysis

All somatic variants that led to a frameshift insertion, frameshift deletion, in-frame insertion, in-frame deletion, missense, nonsense, nonstop, or splice site/region mutation, or a translation start site were considered. For variants identified as missense or nonsense, we required the variant to be considered a likely functional driver using the LiFD tool^18^, which is a two-phase algorithm that pulls from various databases and bioinformatic methods to determine whether a given mutation is likely to be functional. We also considered genes that were significantly mutated in large pediatric cancer and osteosarcoma sequencing studies^4–9, 53^. The final list consisted of 639 genes.

### Evolutionary Analysis

To determine the spatial and temporal dynamics of subclonal diversity within a patient, we first used Treeomics v1.9.2^54^ to derive phylogenies. Treeomics reconstructs the phylogeny of metastatic lesions and maps subclones to their anatomical locations. Treeomics utilizes a Bayesian inference model to account for error-prone sequencing and differing neoplastic cell contents to calculate the likelihood that a specific variant is present or absent. Treeomics then infers a global optimal tree based on mixed-integer linear programming^54^.

The HATCHet^20^ v1.0.1 algorithm was used to infer allele and clone-specific copy-number aberrations, clone proportions, and whole-genome duplications (WGD) for each patient. HATCHet was run using the GATK4-CNV custom pipeline, with Battenberg copy number calls fitted to meet the input requirements for running the tool. Solutions were manually reviewed with the creator of the tool, Simone Zaccaria PhD, to allow for advanced fine-tuning and ensure that the most accurate solutions were selected.

The DeCiFer^17^ v1.1.5 algorithm was used to cluster mutations across all samples for each patient, providing descendant cell fractions (DCF) and cancer cell fractions for each cluster. The copy number input for this algorithm is the output of the HATCHet algorithm. Custom state trees were generated utilizing a maximum copy number between 6-8 for each patient (lower maximum copy number states were selected if the runtime exceeded 48 h). After clustering of mutations, the CALDER^26^ v0.11 algorithm was used to infer evolutionary phylogenies. To run the DeCiFer output through CALDER, the inferred cluster DCF was converted to a read count by multiplying the DCF by 1000 and then dividing by 2 (because CALDER assumes that all mutations are in heterozygous diploid regions). Therefore, if a mutation has 1000 reads and an inferred DCF of 40%, the corresponding input will have 200 variant reads and 800 reference reads. Longitudinal constraints were lifted when analyzing OSCE2 and OSCE3, because all analyzed samples were present at diagnosis.

The Palimpest^23^ algorithm (version = github commit 4795da2) was used to characterize and visualize mutational signatures (using Cosmic SBS/DBS v3.2) at both clone and sample levels. This algorithm also provided information regarding the timing of duplication and loss of heterozygosity events using previously described methods^24^.

### Structural Variant Analysis

Structural variants were annotated using iAnnotateSV software^55^. We used svpluscnv^56^ and ShatterSeek^57^ to identify regions of chromothripsis. The JaBba tool^25^ was used to identify regions with complex rearrangements or amplicon events. The calls from all tools were combined for further downstream analysis.

### Data Visualizations

Oncoprint was created using the CoMut^58^ tool Timescape^59^ (https://github.com/shahcompbio/timescape) and Mapscape^59^ (https://github.com/shahcompbio/mapscape) were used to visualize temporal and spatial clonal evolution. Tableau Desktop (v2021.4) was used to analyze and visualize the data with charts, bars, and line graphs. Ridgeline graphs were created using R utilizing the ggridges package (https://cran.r-project.org/web/packages/ggridges/vignettes/introduction.html). Sankey plots were created using SankeyMATIC (https://sankeymatic.com/build/). Anatomic cartoons were created using BioRender (https://biorender.com/).

### Data Availability

Sequence data will be deposited at the European Genomephenome Archive (EGA), which is hosted by the European Bioinformatics Institute and the Centre for Genomic Regulation. Further information about EGA can be found at https://ega-archive.org and “The European Genome-phenome Archive of human data consented for biomedical research” (http://www.nature.com/ng/journal/v47/n7/full/ng.3312.html).

