## Supplementary Figures for "Subclonal somatic copy number alterations emerge and dominate in recurrent osteosarcoma"

### Supplementary Figure 1

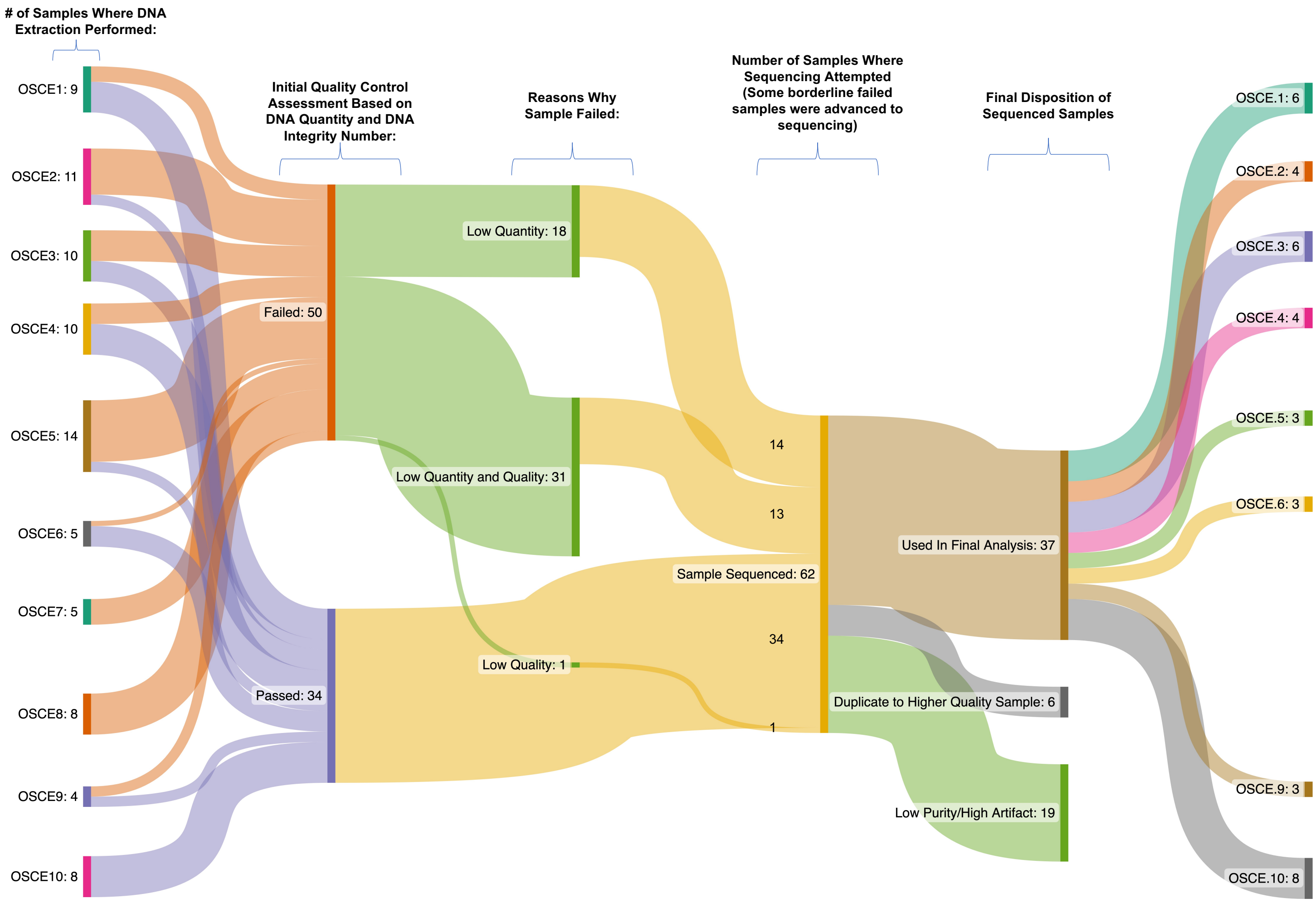


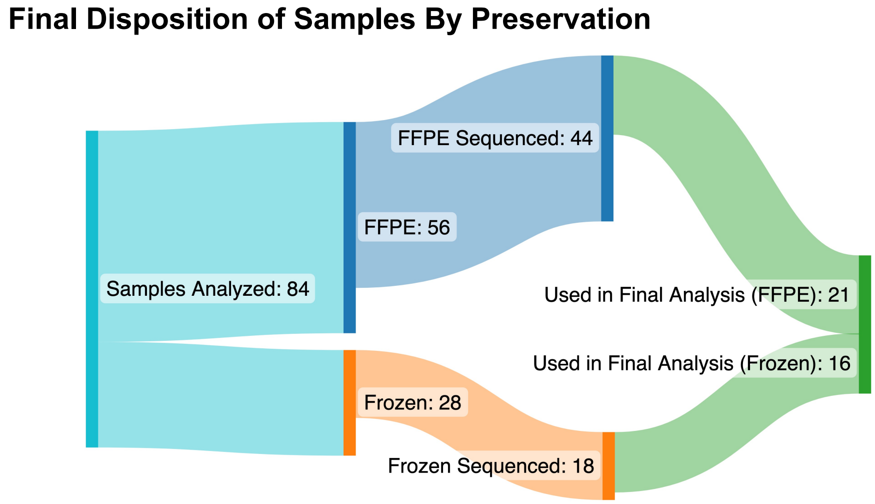

**Supplementary Figure 1.** Sankey plots of sample disposition. The upper plot shows the number of samples that had DNA extracted for each patient, then flows to whether those sample passed our Integrated Genomics Operation (IGO) standards to proceed with whole genome sequencing, followed by why a sample failed initial quality control. Some samples that failed initial quality control were advanced to whole genome sequencing due to the rare nature of these samples. Of the 62 samples that were sequenced 37 were used in the final analysis, with reasons why a samples was discarded at this point shown on the plot. At the bottom of the plot is the final disposition by sample preservation. Starting with 84 samples, 44 FFPE samples were sequenced, while 18 frozen samples were sequenced with 21 FFPE and 16 Frozen samples selected for final analysis.

**Supplementary Figure 2**

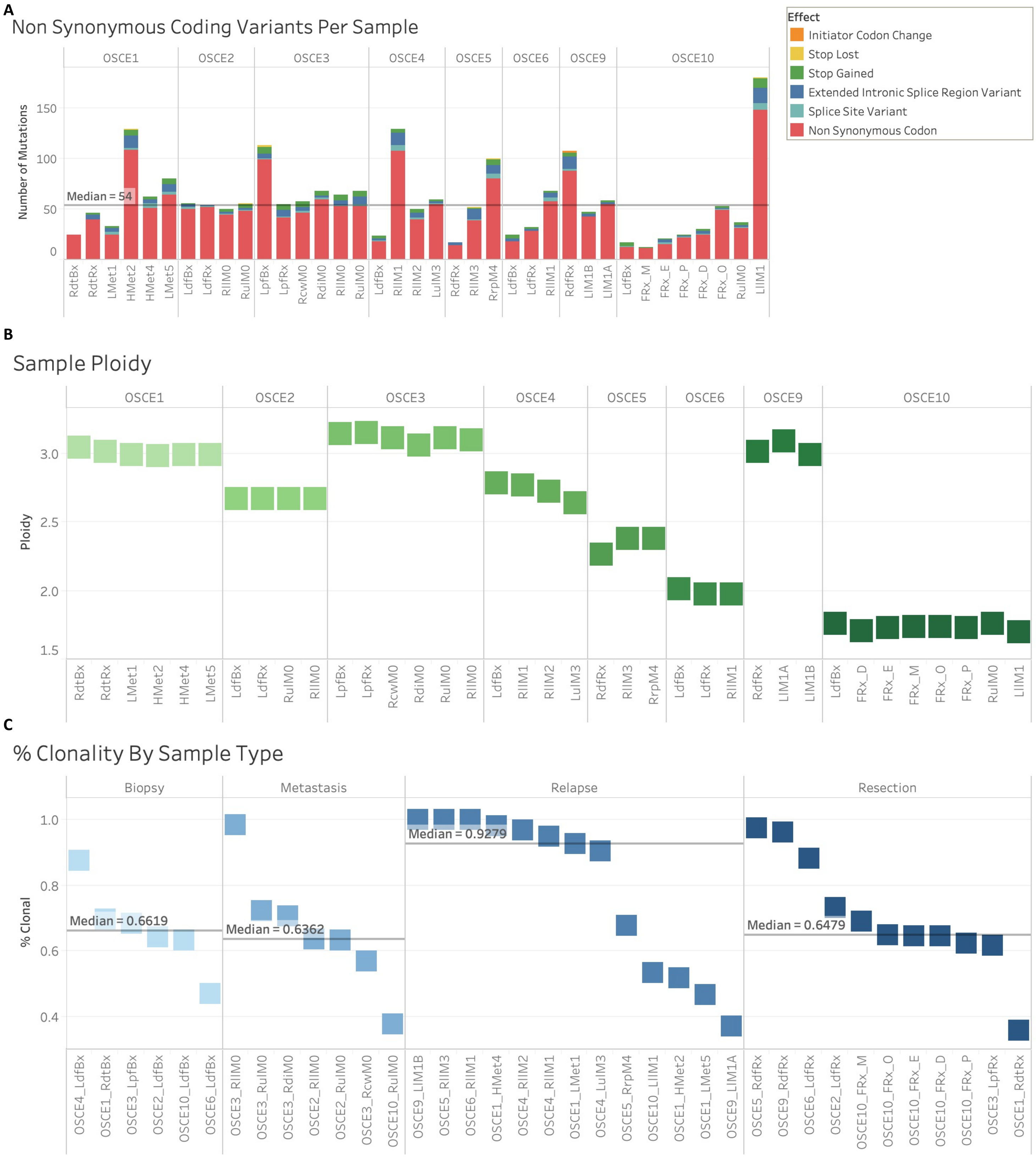


12

**Supplementary Figure 2.** Extended genomic characteristics of the samples in our cohort. **A,** Stacked bar plot of the non-synonymous coding variants for each sample. Colors represent the different type of non-synonymous variants and their bar plot size is based on the number of mutations. **B,** Sample ploidy for each patient, samples are ordered from earliest on left to most recent on right. **C,** Plot of the percentage of mutations considered to be clonal. Mutations were considered clonal based on a descendant cell fraction ≥ 90% (analogous to cancer cell fraction). Samples are ordered from earliest on left to most recent on right.

### Supplementary Figure 3

A

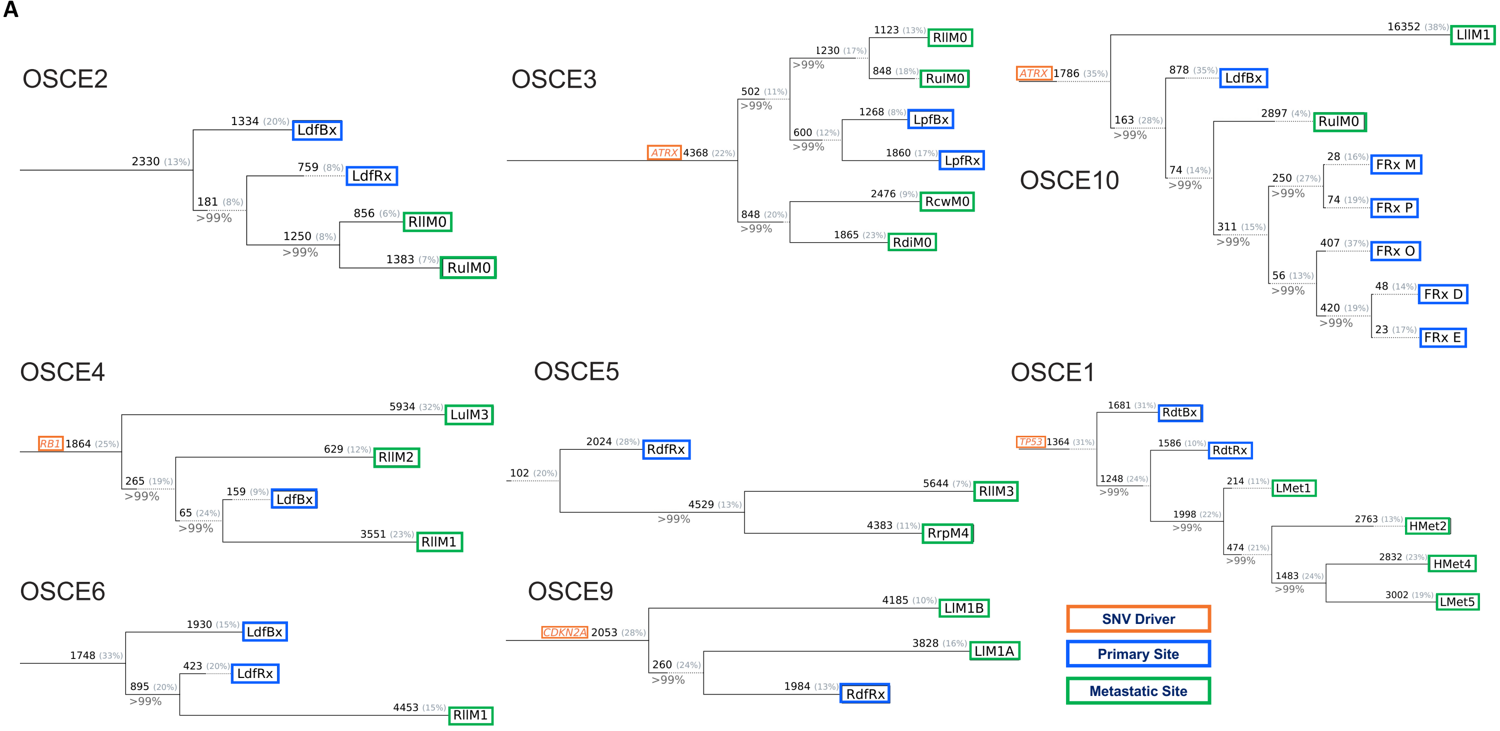


B

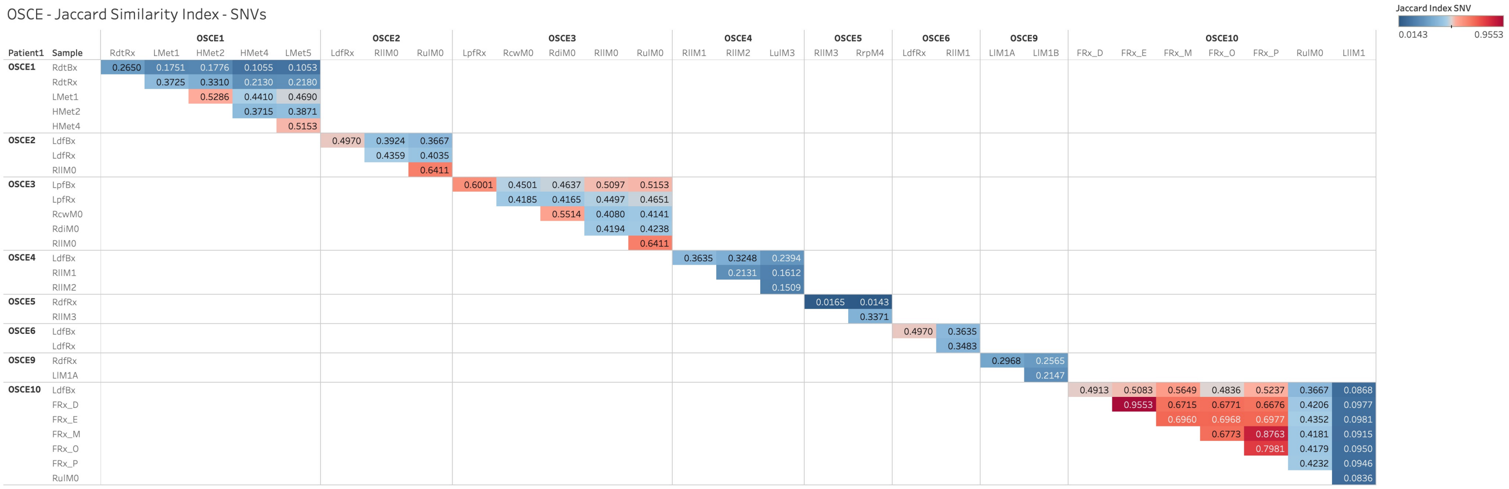


**Supplementary Figure 3.** Sample based phylogenies as determined by the Treeomics algorithm. **A,** In the five patients (OSCE1, OSCE3, OSCE4, OSCE9, OSCE10) who were found to have likely functional driver SNVs, these mutations were found to be truncal in each phylogeny.

Seeding patterns across the cohort was heterogenous. For 4 patients (OSCE1, OSCE2, OSCE5, OSCE6), the primary tumor biopsy or resection formed the outgroup in the phylogeny (all metastatic lineages share a common ancestor that is more recent than their most recent common ancestor with primary tumor), indicating that recurrent disease developed from a single resistant clonal population (monophyletic seeding). For the remaining 4 patients (OSCE3, OSCE4, OSCE9, OSCE10), the recurrent disease was inferred to be seeded by multiple ancestral clones (polyphyletic seeding). **B,** To objectively assess relatedness among primary and metastatic/recurrent disease, we calculated pairwise Jaccard similarity coefficients for all samples within a patient, observing similar patterns to what Sakamoto[1] et al. described, with monophyletic recurrences more distant (lower Jaccard similarity coefficient) than polyphyletic recurrences, which were more related to the primary tumor.

### Supplementary Figure 4

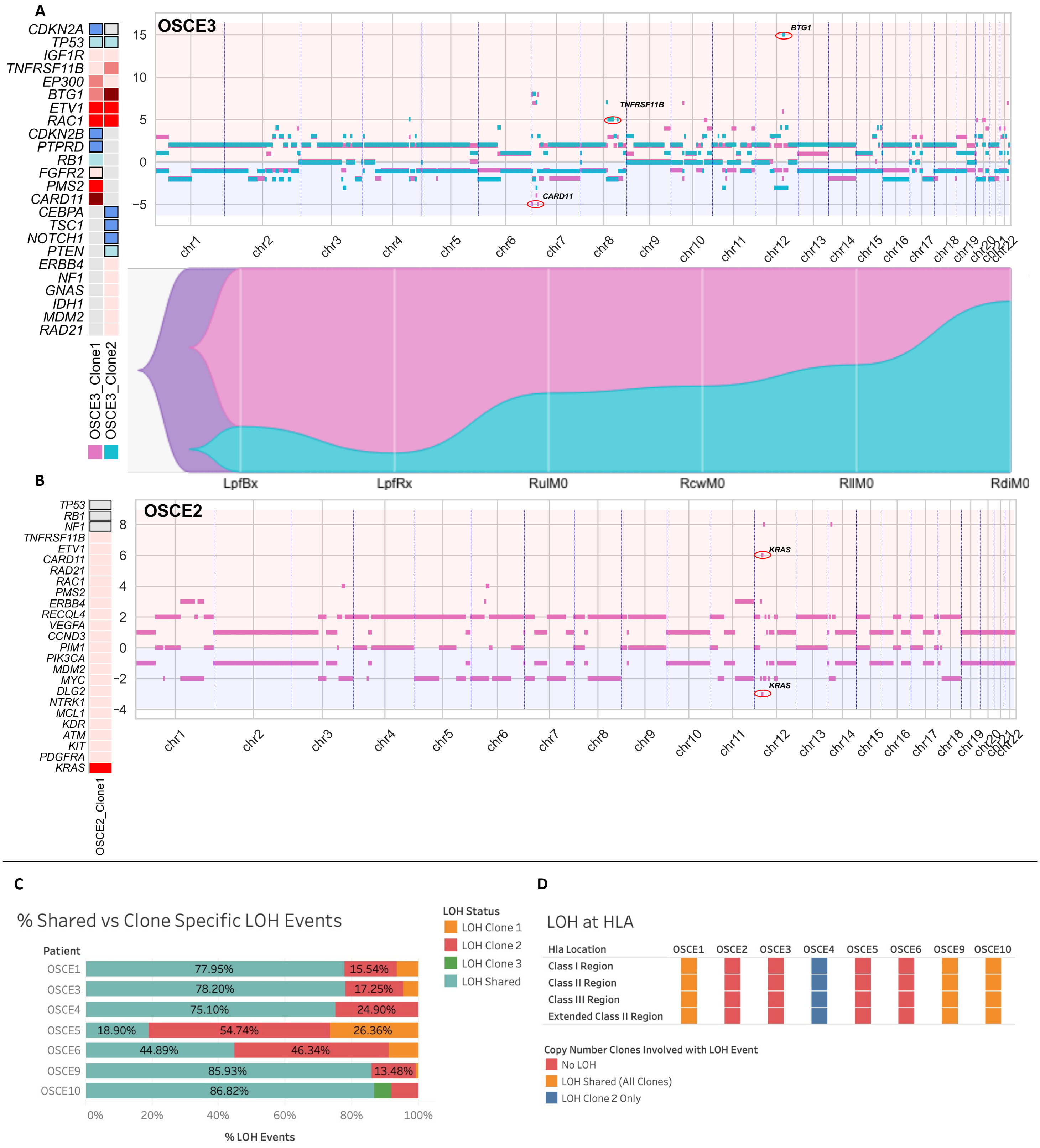


35

**Supplementary Figure 4.** Extended analysis of copy number alterations, clones, and clone proportions. **A,** and **B,** For each patient there is a panel of three figures (except OSCE2 as there was only 1 copy number clone). The figure on the left is an oncoprint featuring clone specific copy number alterations in recurrently altered genes of interest in osteosarcoma. The top figure is a plot of allele specific copy number alterations for each clone with significant events for each clone circled and highlighted. Clone 1 is the magenta clone and clone 2 is the teal clone. The major allele is plotted above 0 and the minor allele is plotted below 0. The bottom figure in each panel is a TimeScape plot of the prevalence of each clone at different timepoints throughout a patient's disease course. **C,** Stacked bar chart of showing the percentage of shared vs private LOH events for the copy number clones found in each patient. **D,** LOH status at the HLA locus for the copy number clones found in each patient.

**Supplementary Figure 5**

**A**

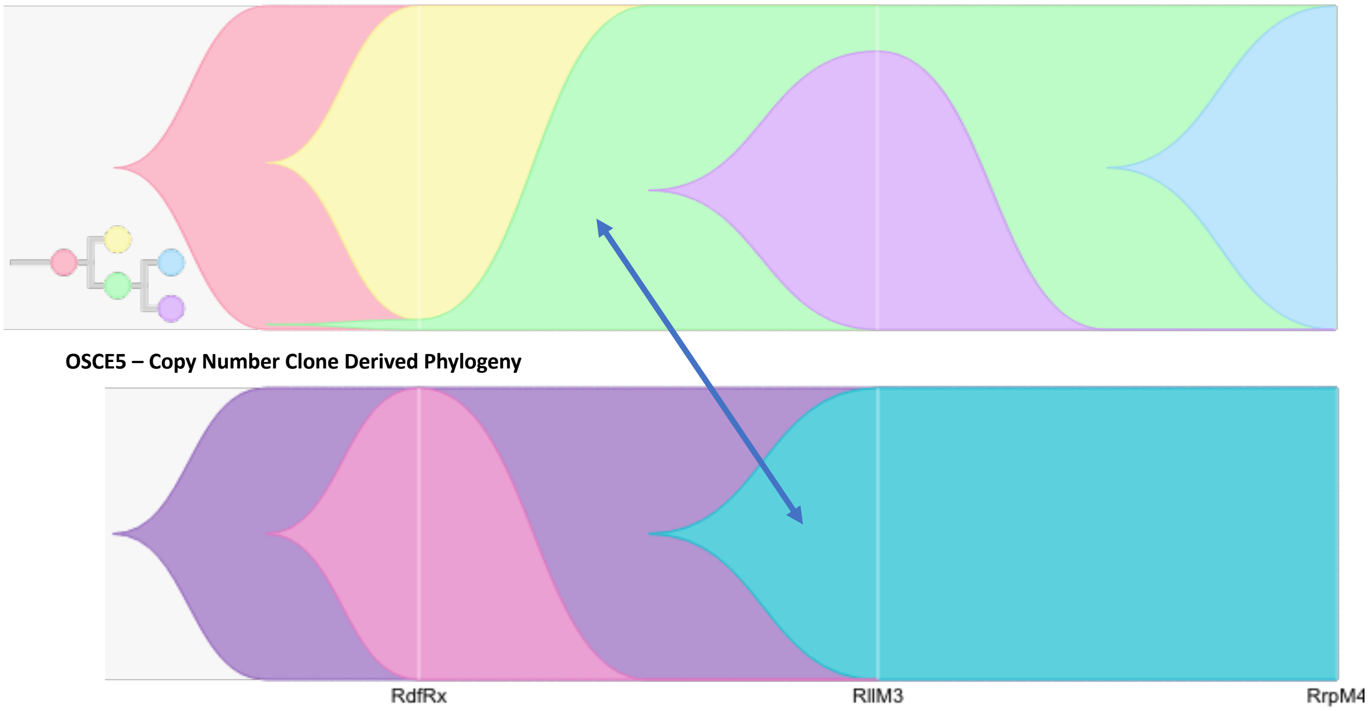


**B**

### MYC Amplification Characterizes Resistant Copy Number Clone

| Patient | Classification | Clone 1 | Clone 2 | Clone 3 |
|---|---|---|---|---|
| OSCE1 | Low Amp | | + | |
| OSCE4 | Gain | | + | |
| OSCE5 | Gain | + | | |
| | High Amp | | + | |
| OSCE6 | Gain | + | | |
| | Low Amp | | + | |
| OSCE9 | Gain | + | | |
| | Mid Amp | | + | |
| OSCE10 | Low Amp | + | + | |
| | High Amp | | | + |

Classification
- Gain
- Low Amp
- Mid Amp
- High Amp

**Supplementary Figure 5. A,** Comparison of SNV derived phylogeny vs copy number derived phylogeny. The teal copy number clone emerges and is a major clone for RllM3, similar to the green SNV based clone. Due to the fact it is easier to track subclonal populations of SNVs, we see that the green clone can be rooted in the primary sample and then follows a similar clonal trajectory as the teal clone. The most likely explanation is that the teal clone was likely present at diagnosis as well but could not be detected due to the limitations of detecting subclonal copy number populations. **B,** Plot of the type of *MYC* gains/amplifications that characterize each clone.

### Supplementary Figure 6

### Duplication Timing in Patients with Ploidy ≥ 2.5

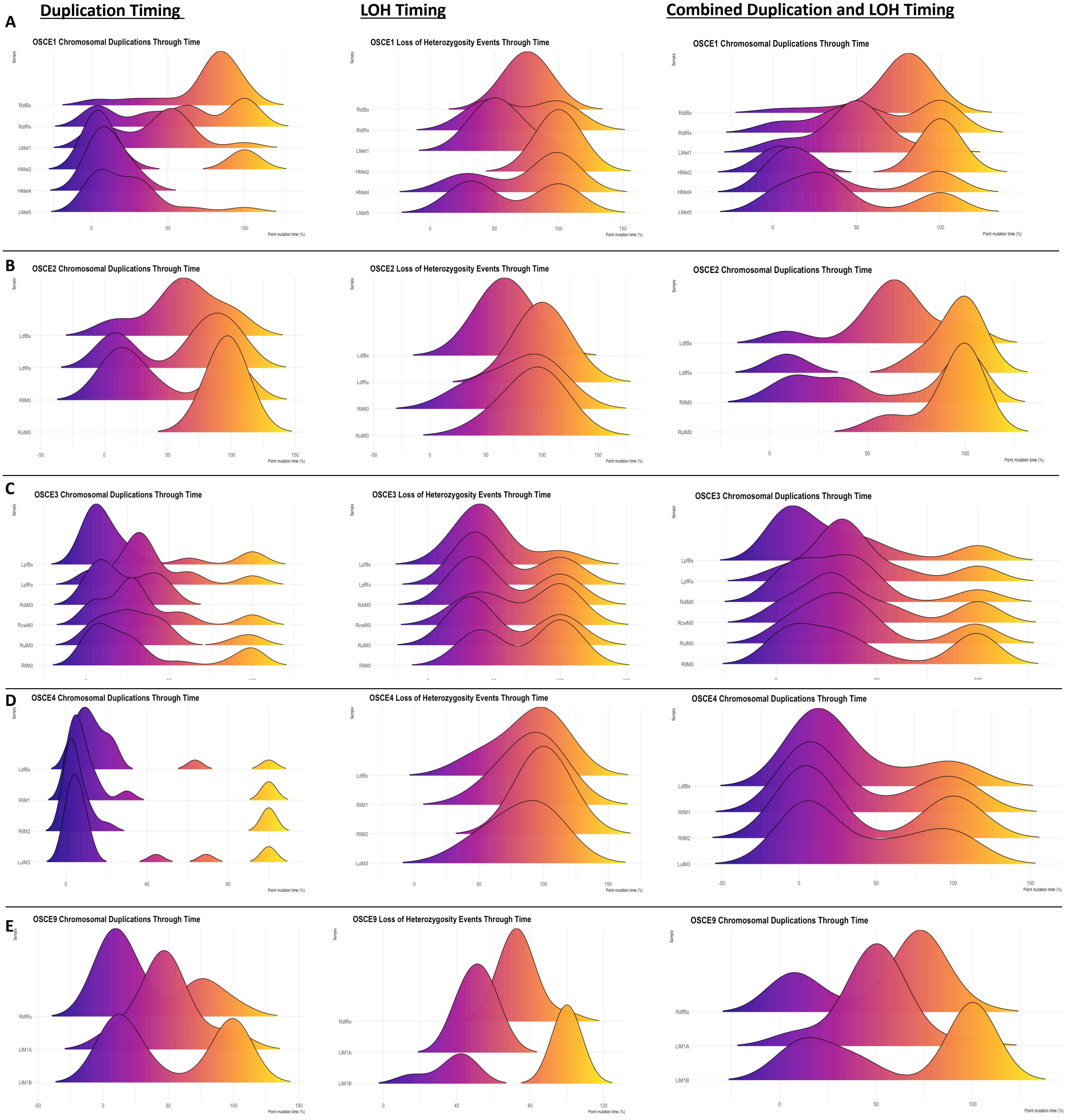

**Supplementary Figure 6**. Duplication timing in patients with ploidy ≥ 2.5. **A, B, C, D, and E,** Duplication, LOH, and combined duplication and LOH ridgeline plots are shown for each selected patient. Each row represents a different patient. Samples for each patient are shown from earliest collected sample at the top to the most recent sample at the bottom of each plot.

### Supplementary Figure 7

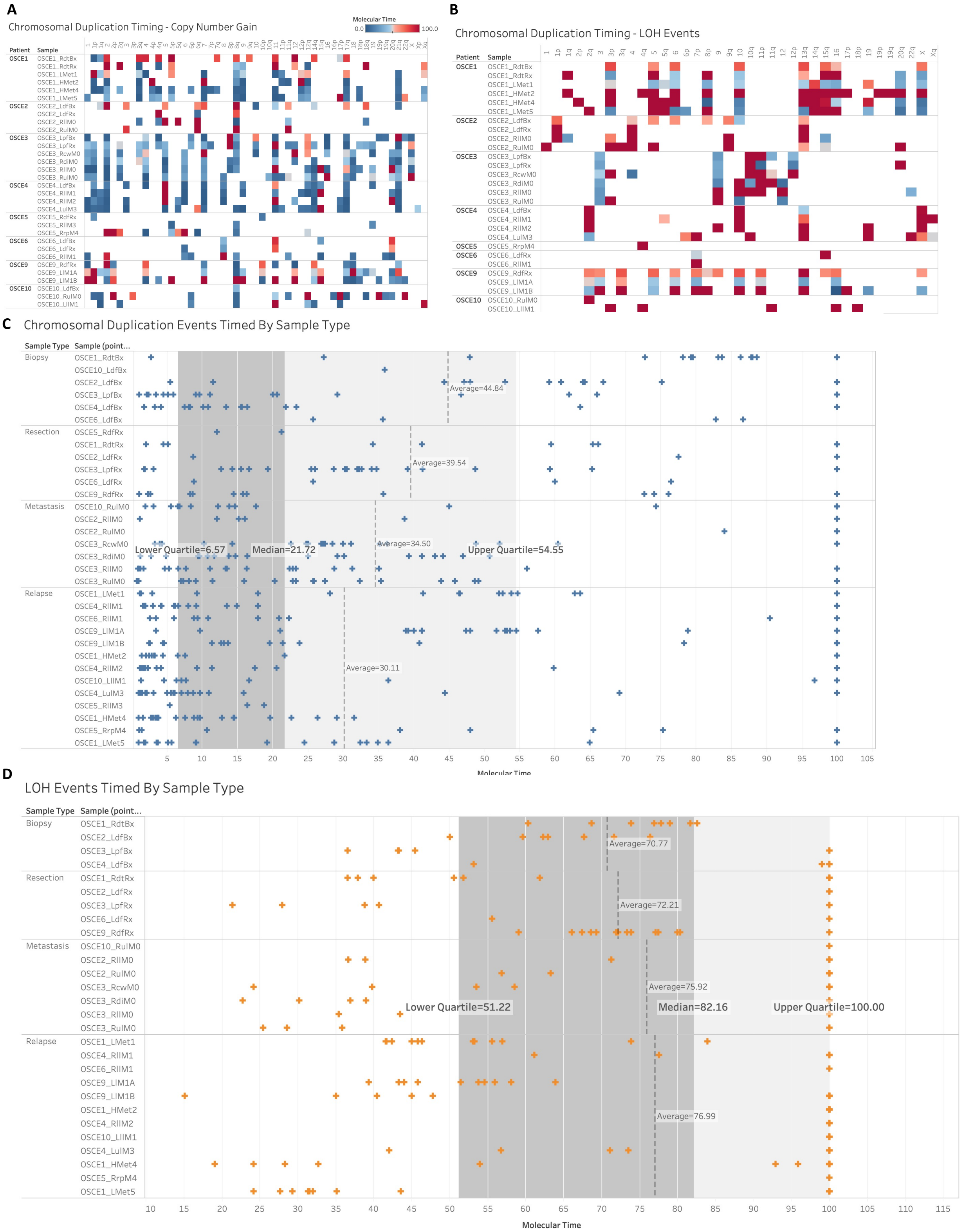


61

**Supplementary Figure 7.** Extended duplication timing analysis. **A,** Heatmap of chromosomal duplication timing by arm involved. **B,** Heatmap of LOH timing by arm involved. **C,** Plot of chromosomal duplication events plotted along x-axis of molecular time and arranged by sample type. Average duplication time for each type of sample is represented by the vertical dotted line. The lower quartile (dark gray), median, and upper quartile (light gray) are shaded for the entire cohort. **D,** Plot of LOH events plotted along x-axis of molecular time and arranged by sample type. Average LOH time for each type of sample is represented by the vertical dotted line. The lower quartile (dark gray), median, and upper quartile (light gray) are shaded for the entire cohort.

**Supplementary Figure 8**

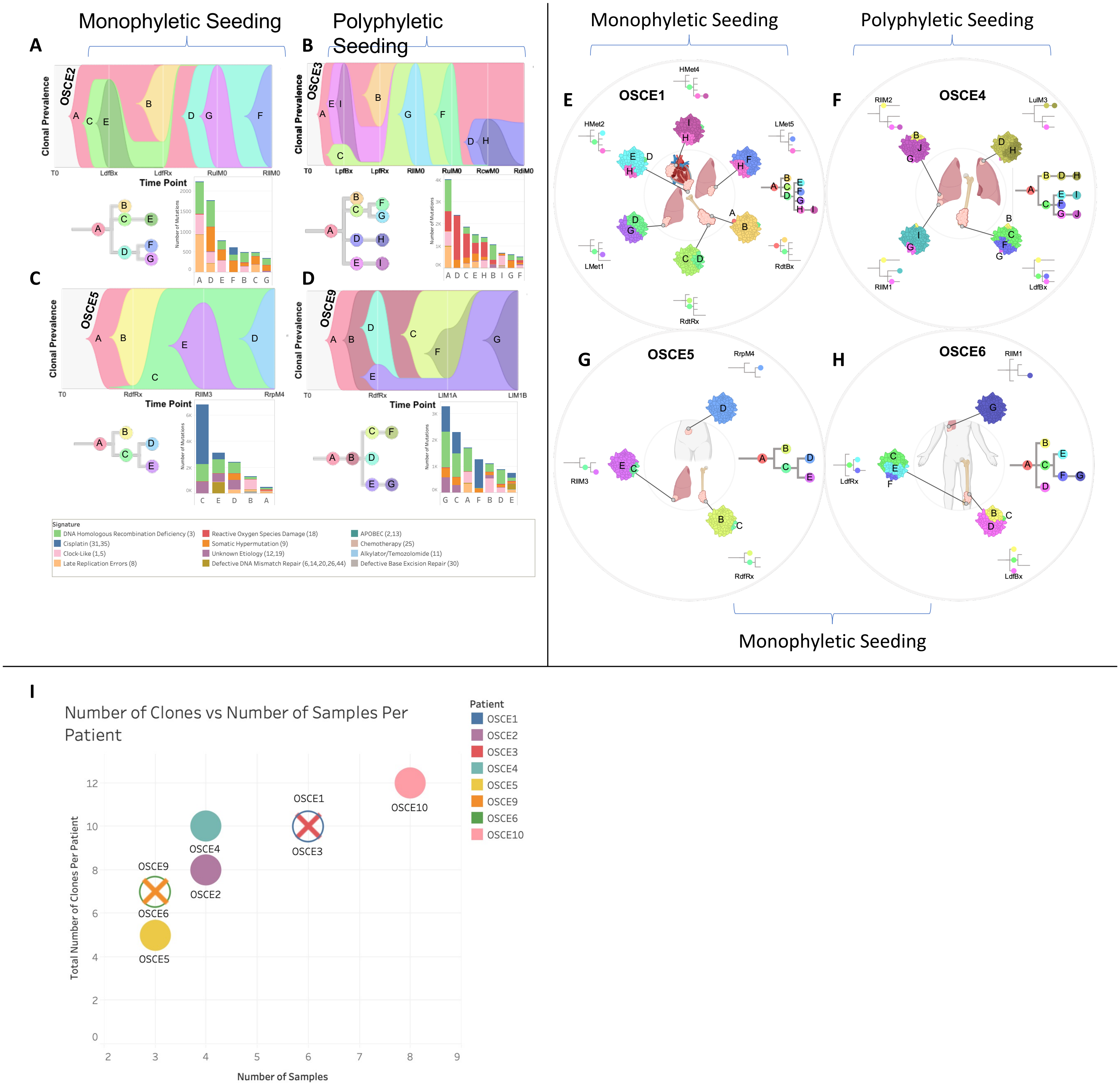

**Supplementary Figure 8.** SNV based phylogenies. **A, B, C, and D,** Upper figure in each panel is a TimeScape plot of the inferred evolutionary phylogeny, highlighting clonal proportions over time. The prevalence of different clones is shown over time on the vertical axis, with the different clones represented by different colors. The horizontal axis represents the timepoints, which are represented by gray lines. The evolutionary relationships between the clones are shown on the phylogenetic tree and in the TimeScape layout. The bottom right of each panel is a stacked bar plot of the total number of mutations assigned to each clone. Colors represent total number of mutations attributed to each mutational signature with color legend at bottom of figure. Patient level metastatic seeding patterns are denoted by the brackets at the top of the page. **E, F, G, and H,** Spatially and in some cases temporally distinct samples are indicated on the anatomic sites from where the sample originated. The colors represent different clones, and the phylogenetic trees show the evolutionary relationships between these clones. The prevalence of each clone at a particular site is proportional to the colored area of the cellular aggregate representation. **I,** The number of clones are plotted on the y-axis and the number of samples for each patient are plotted on the x-axis. Colors represent the different samples.

### Supplementary Figure 9

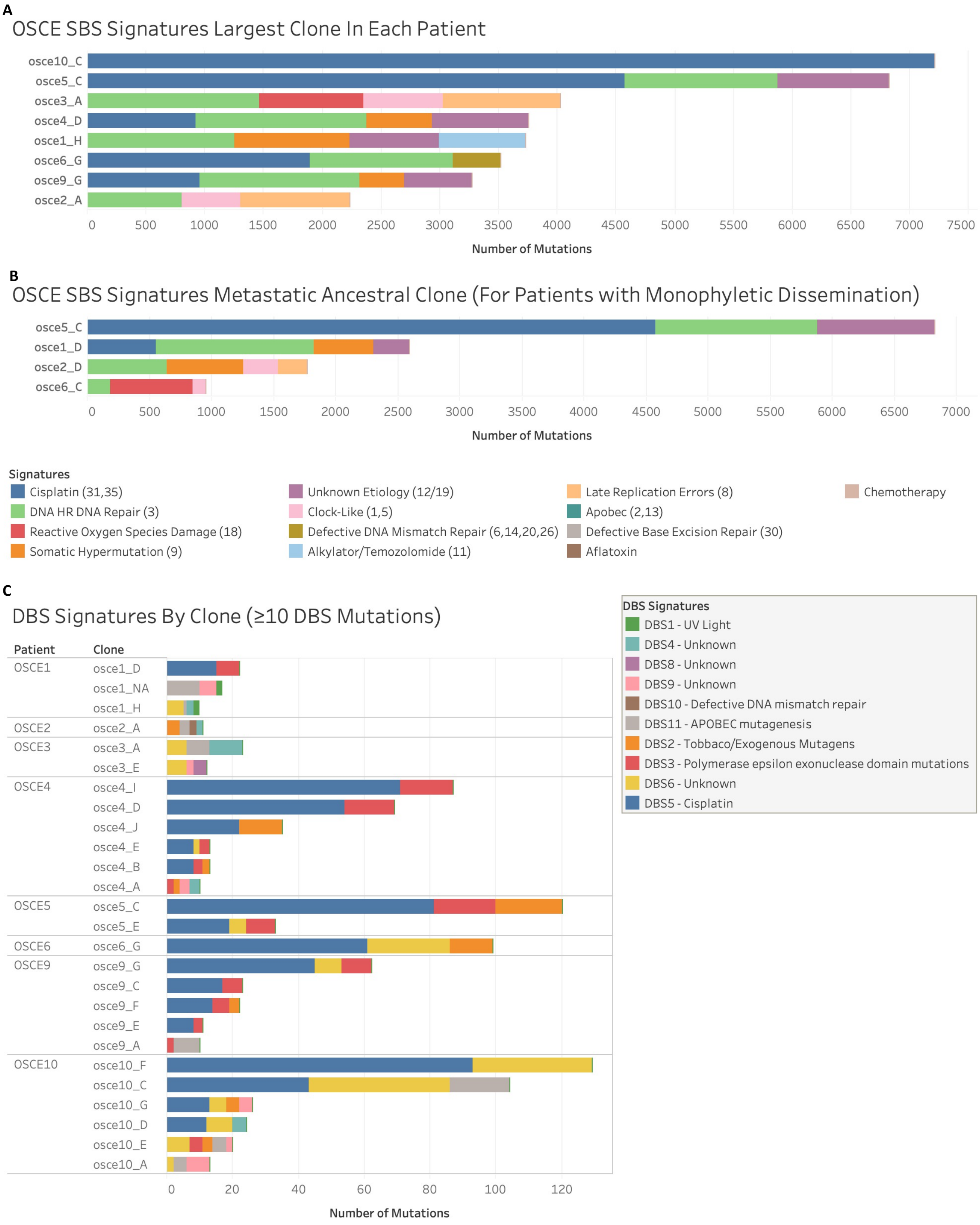

**Supplementary Figure 9.** SBS and DBS mutational signatures across different clones. **A,** Mutational signature breakdown by number of total mutations attributed to each signature for the largest clone by total number of mutations for each patient. Colors represent different signatures. **B,** For patients with monophyletic (when all metastatic clones are derived from a recent common ancestor) dissemination, the mutational signature breakdown by number of total mutations attributed to each signature for the ancestral clone is shown. Colors represent different signatures. **C,** DBS signature breakdown by number of total mutations attributed to each signature for any clone with ≥10 DBS mutations. Colors represent different signatures.

### Supplementary Figure 10

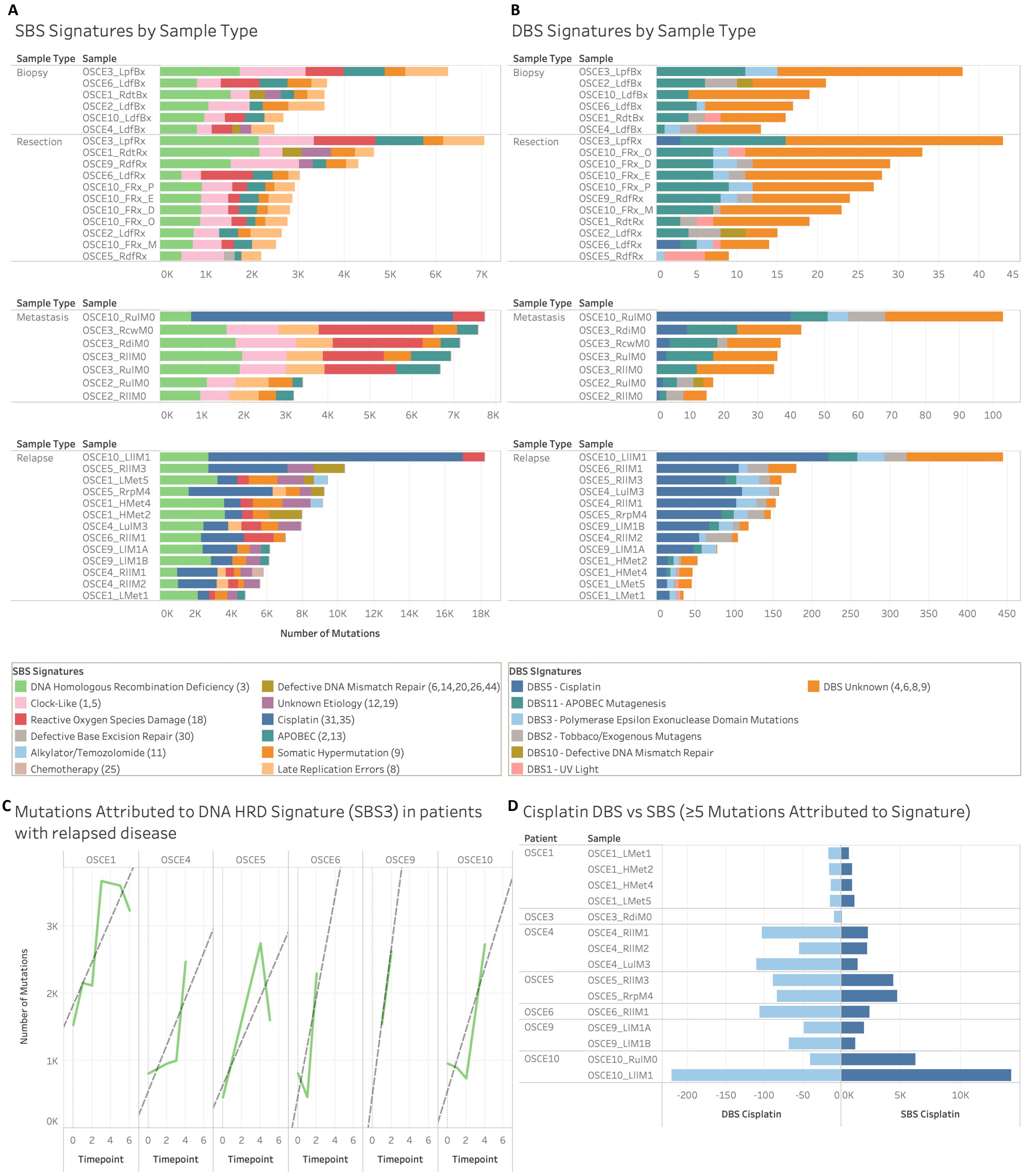

**Supplementary Figure 10.** SBS and DBS mutational signatures at the sample level with temporal trends of the HRD-related SBS3 and cisplatin signatures. **A,** SBS mutational signature breakdown by number of total mutations attributed to each signature for each sample organized by sample type. Colors represent different signatures. **B,** DBS mutational signature breakdown by number of total mutations attributed to each signature for each sample organized by sample type. Colors represent different signatures. **C,** Temporal trend of total number of mutations attributed to HRD-related SBS3 at sequential timepoints for each patient. **D,** Number of mutations attributed to SBS and DBS signatures in samples with ≥5 mutations attributed to these signatures. DBS Signatures are represented by the light blue and are plotted along the x-axis, with increasing number of mutations moving right to left. SBS signatures are represented by the dark blue and are plotted along the x-axis, with increasing number of mutations moving left to right. Note the difference in scales along the x-axis for DBS and SBS respectively.
